# D-lactate enhances inflammatory and cytotoxic functions of Th1 cells through stereospecific control of mitochondrial ROS

**DOI:** 10.1101/2025.04.03.647060

**Authors:** Adrián Madrigal-Avilés, Valerie Plajer, Paulo R. Viana Mendonça, Sebastian Serve, Leticia Soriano-Baguet, Dirk Brenner, Maria Dzamukova, Mir-Farzin Mashreghi, Max Löhning

**Author notes:** Correspondence: Max Löhning; Adrián Madrigal-Avilés.

## Abstract

CD4+ T lymphocytes with cytotoxic activity are an understudied T helper (Th) cell population with the capacity to orchestrate immune responses and kill target cells in a TCR-restricted manner. Particularly, the mechanisms that promote the differentiation of cytotoxic Th cells remain elusive. Here, by supplementing Th1 cells with different lactate stereoisomers, we identify a previously unrecognized stereospecific adaptation that reinforces cytotoxic and proinflammatory programs. Although both D- and L-lactate impacted histone acetylation and metabolic activity, only D-lactate increased IFN-γ and TNF-α production and most potently promoted cytotoxic function. D-lactate also selectively enhanced mTORC1 and ERK1/ERK2 signaling, although these changes were not sufficient to fully explain the distinctive phenotype. Instead, the decisive feature of D-lactate supplementation was a reduction in mitochondrial reactive oxygen species (ROS), associated with electron transport chain (ETC) remodeling and enhanced glyoxalase-dependent buffering. Together, these findings identify a stereospecific lactate redox axis promoting cytotoxic Th1 cell differentiation.

## Introduction

Upon activation, naïve CD4^+^ T cells can differentiate into various subsets depending on the cytokine environment. Viral infections, intracellular pathogens and tumors promote Th1 differentiation. Interferon-γ (IFN-γ) and interleukin-12 (IL-12) activate signal transducer and activator of transcription 1 (STAT1) and STAT4, respectively, inducing T-bet, tumor necrosis factor-α (TNF-α) and IFN-γ. A fraction of Th1 cells can further differentiate into cytotoxic CD4□ T cells that kill target cells through granzyme B (GZMB) and perforin-1 (Perf-1)^1,2^.

Beyond cytokine and transcription factor control, metabolic adaptations also govern Th differentiation. Naïve T cells remain quiescent largely through basal mitochondrial oxidative phosphorylation (OXPHOS), supported by the tricarboxylic acid (TCA) cycle and electron transport chain (ETC), which generates the membrane potential driving ATP production. Perturbations in mitochondrial electron transport generate mitochondrial reactive oxygen species (mitoROS), which can be converted into signaling molecules such as H O that affect transcriptional programs and cytokine production^3–5^. Their detoxification relies on antioxidant systems centered on glutathione (GSH). In CD4□ T cells, ROS shape effector function in a subset-dependent manner, favoring IL-4 production in Th2 cells while negatively affecting IFN-γ and IL-10 production in Th1 and Treg cells, respectively^6–8^. However, the metabolic adaptations underlying acquisition of cytotoxic features in CD4□ T cells remain largely unknown.

Upon antigen recognition, T cells shift toward aerobic glycolysis to support proliferation^9^. This metabolic rewiring is orchestrated by mTORC1, which promotes glycolytic programs and protein synthesis^10,11^. As a result, pyruvate is preferentially converted into lactate rather than fully oxidized through mitochondrial OXPHOS.

Lactate exists as two enantiomers, D- and L-lactate, which differ in the configuration of their asymmetric carbon^12^. Their metabolism is stereospecific: L-lactate is interconverted with pyruvate by cytosolic LDHA and LDHB, whereas D-lactate is metabolized by mitochondrial LDHD^13,14^. In higher organisms, L-lactate is the predominant glycolytic end-product. Glycolytic flux also generates methylglyoxal (MG), a highly reactive dicarbonyl detoxified into D-lactate by the GSH-dependent glyoxalase system^15–17^. In immune cells, MG dampens lymphocyte activation and cytokine production.^18,19^.

Beyond MG detoxification, D-lactate is linked to metabolically active effector immune cells and is generated by commensal bacteria through carbohydrate fermentation. Barrier disruption in inflammatory bowel disease (IBD) allows microbial metabolites to disseminate into the circulation, and D-lactate levels are used to track disease activity^20,21^. IBD is also characterized by IFN-γ–producing cytotoxic CD4□ T cells ^22,23^, which reside at sites of barrier disruption and may encounter high local D-lactate. Whereas systemic D-lactate levels remain micromolar, bacterial fermentation of malabsorbed carbohydrates^24^ can raise luminal D-lactate to 110 mmol/L in the dysbiotic colon^25^. Whether D-lactate actively shapes these effector states remains unexplored.

Additionally, lactate can influence gene expression through chromatin remodeling mechanisms^26^. Lactate metabolism can increase acetyl-CoA availability for histone acetylation^27^, both lactate enantiomers have been reported to exert mild histone deacetylase (HDAC) inhibitory activity^26^ and intracellular lactate accumulation can also promote lactylation, a recently described post-translational modification linked to transcriptional and metabolic regulation^28,29^. Together, these mechanisms give lactate the potential to reshape the regulatory landscape underlying immune effector differentiation^28^. At the same time, excessive L-lactate accumulation has been shown to impair Th1 differentiation and IFN-γ production^28,29^.

While several studies have addressed the role of L-lactate in CD4^+^ T cell biology^6,27,30^, very little is known about the effects of D-lactate in immunity^31,32^. In this study, we compared the impact of D- and L-lactate supplementation on Th1 differentiation, integrating inflammatory, cytotoxic, metabolic, redox, and chromatin-associated readouts, and reveal that the two enantiomers drive distinct effector states through stereospecific control of mitochondrial ROS.

## Results

### D-lactate enhances proinflammatory cytokine production and cytotoxic function in Th1 cells

To determine how lactate stereoisomers influence Th1 differentiation and effector function, we cultured sorted mouse naïve CD4□ T cells for 5 days under Th1-polarizing conditions with or without 20 mM Na-D-lactate or Na-L-lactate (hereafter D- and L-lactate). D-lactate increased the frequency of IFN-γ-producing cells and the per-cell abundance of IFN-γ and TNF-α (MFI), whereas L-lactate reduced both proinflammatory cytokines relative to control (Figures 1A and 1B), indicating that the two stereoisomers induce opposite inflammatory phenotypes.

**Figure 1.**
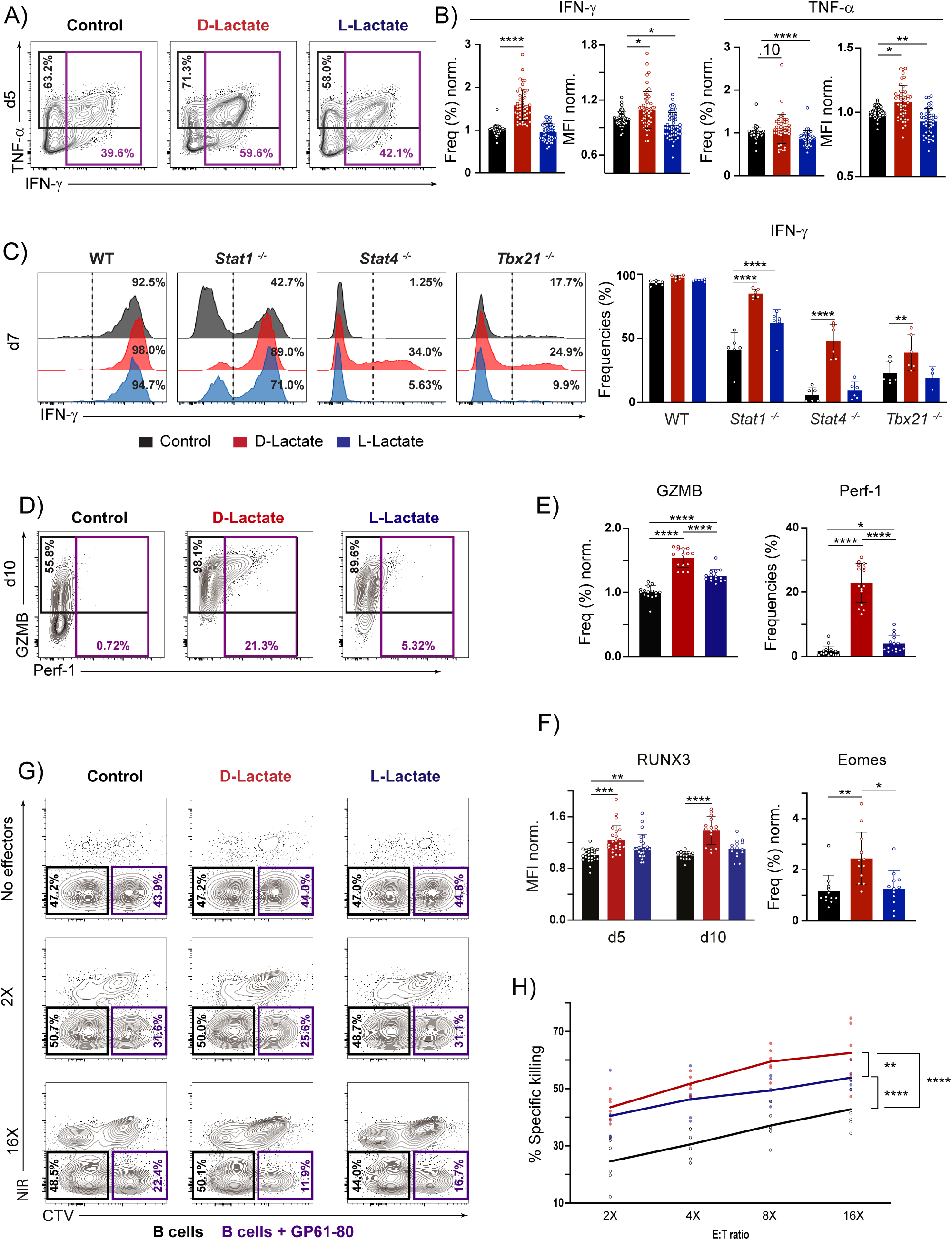
**D-lactate increases IFN-γ production independently of STAT1, STAT4 and T-bet while inducing strong cytotoxic features.** Naïve CD4□ T cells sorted from WT, SMARTA, STAT1-, STAT4- or T-bet-deficient mice were cultured under Th1-polarizing conditions ± lactate as indicated. Analyses on day 5 (A, B, F–H), day 7 (C) or day 10 (D–F). (A) Representative contour plots of IFN-γ and TNF-α. (B) Quantification of (A), frequencies and MFI (n=51 IFN-γ; n=45 TNF-α). (C) Representative histograms (left) and quantification (right) of IFN-γ (n=6). (D) Representative contour plots of GZMB and Perf-1. (E) Quantification of (D) (n=15). (F) Intracellular RUNX3 (left) and EOMES (right) (RUNX3, n=24 (d5) and 15 (d10); EOMES, n=12). (G) Representative contour plots of living B cells after the killing assay at the indicated effector:target (E:T) ratios. (H) Quantification of (G) as area under the curve (AUC) of specific killing across all E:T ratios (n=6). Mixed-effects with Dunnett’s (C), One-way ANOVA with Tukey’s post-tests (E, F (Eomes), H), Kruskal-Wallis with Dunn’s post-tests (B) or Two-way ANOVA with Dunnett’s post-tests (F (RUNX3)).

Because Th1 differentiation is classically coordinated by the synergistic actions of STAT1, STAT4 and T-bet, which together promote IFN-γ transcription ^33^, we next asked whether D-lactate acted through this canonical regulatory axis. As expected, deficiency in *Stat1*, *Stat4* or *Tbx21* dampened overall IFN-γ production, but none of these perturbations abolished the ability of D-lactate to upregulate IFN-γ (Figure 1C). In fact, the effect was stronger than in WT cells, indicating an alternative mechanism to classical Th1-associated factors.

Because cytotoxic CD4□ T cells arise from terminally differentiated Th1 cells during sustained viral infections^34^, we next investigated whether lactate supplementation could influence the acquisition of cytotoxic traits, submitting Th1 cells to an additional round of stimulation under Th1-skewing conditions and analyzing them at day 10. Both stereoisomers increased the proportion of GZMB and Perf-1 cells relative to control, but D-lactate elicited the strongest effect (Figures 1D and 1E). Consistent with this phenotypic shift, expression of cytotoxicity-related transcription factors, including EOMES and RUNX3, was also altered, with D-lactate showing the most pronounced response and L-lactate a modest effect on RUNX3 (Figure 1F).

We next asked whether these changes translated into enhanced antigen-specific cytotoxic function. We therefore used SMARTA transgenic CD4□ T cells, specific for the LCMV GP61-80 peptide^35^ presented by murine MHC-II I-A^b^ molecules^36^. *In vitro* killing assays showed that lactate supplementation augmented Th1 cytotoxicity after only 5 days of differentiation, with D-lactate exhibiting the highest rates at all effector-to-target ratios tested (Figures 1G and 1H). Together, both stereoisomers promote cytotoxic features in Th1 cells, but D-lactate exerts a significantly stronger effect, coupling enhanced cytotoxic differentiation to increased proinflammatory cytokine production.

### Lactate stereoisomers share rapid histone hyperacetylation, whereas only D-lactate remodels chromatin accessibility

We next measured histone 3 (H3) acetylation at K9 and K27, associated with active promoters and enhancer/superenhancer activity^37,38^, respectively, comparing prolonged lactate treatment (5 days) with acute 3-hour exposure in day 5 control-differentiated cells during PMA/ionomycin restimulation. Both stereoisomers significantly increased H3K9 and K27 acetylation, consistent with previous reports, and acute exposure induced hyperacetylation at both residues, with Ac(K27)-H3 levels slightly higher than after prolonged exposure (Figures 2A and 2B). However, despite comparable acetylation between stereoisomers and exposure durations, only prolonged D-lactate treatment increased *Ifng* mRNA levels (Figure 2C). To test whether histone acetylation alone induced the Th1 effector program, we supplemented control samples with the HDAC inhibitor SAHA. Unexpectedly, SAHA not only failed to increase TNF-α and IFN-γ expression but instead reduced production of both cytokines (Figure 2D). Histone hyperacetylation is therefore an early response to lactate exposure, but is not sufficient on its own to drive effector cytokine production.

**Figure 2.**
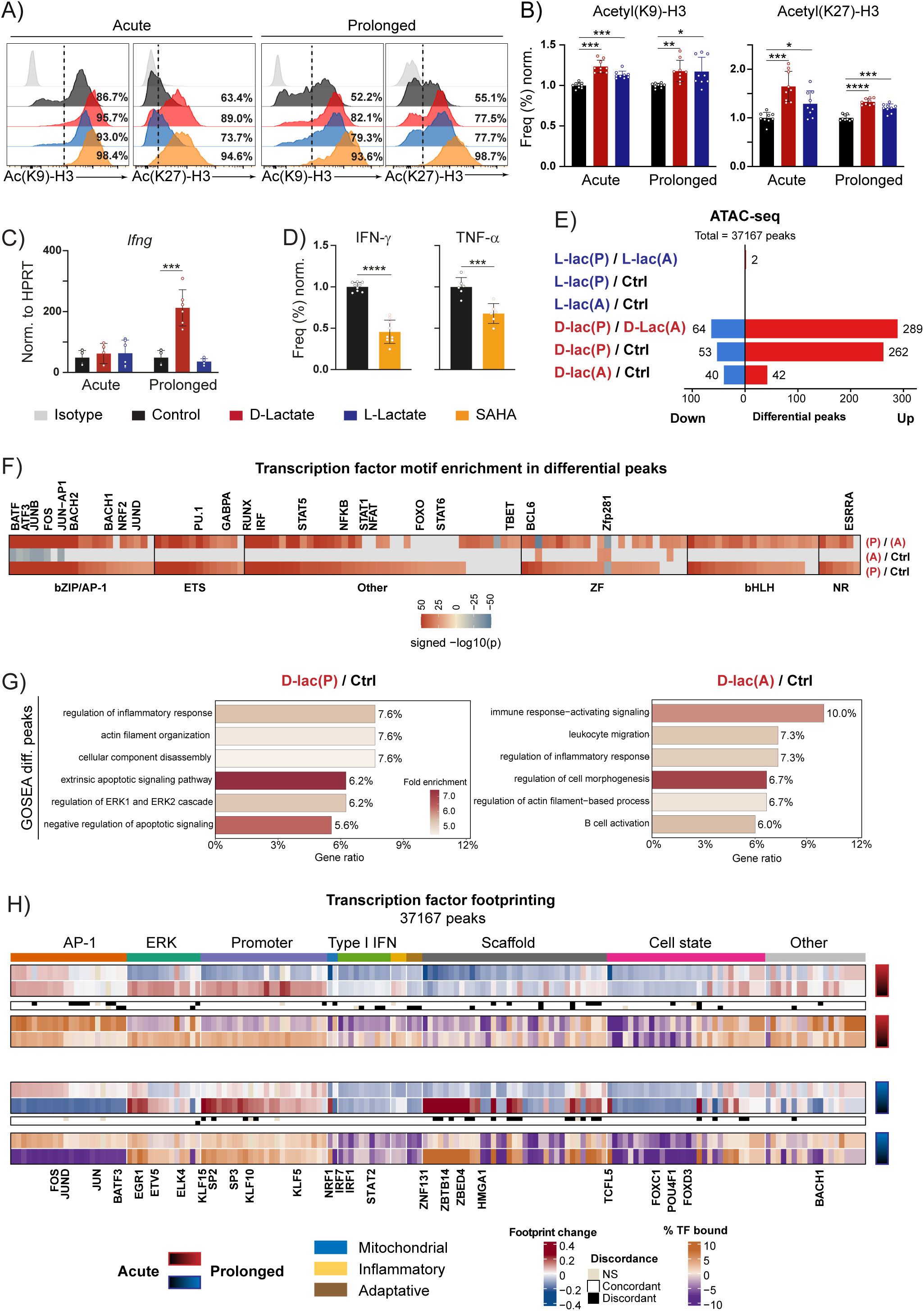
**Early histone acetylation and chromatin remodeling induced by lactate are insufficient to drive the Th1 effector program.** (A–H) Naïve WT CD4□ T cells were cultured for 5 days under Th1-polarizing conditions ± lactate and restimulated with PMA/ionomycin; alternatively, control-differentiated cells were acutely supplemented during restimulation with D- or L-lactate (20 mM) or SAHA (10 μM). (A) Representative histograms of Acetyl(K9)-H3 and Acetyl(K27)-H3. (B) Quantification of (A) (n=9). (C) qRT-PCR of *Ifng* under acute and prolonged lactate treatment (n=6). (D) IFN-γ and TNF-α production after acute SAHA treatment during restimulation (n=9 IFN-γ; n=6 TNF-α). (E) ATAC-seq differential peaks per contrast: prolonged (P) and acute (A). (F) HOMER TF motif enrichment within differential peaks (P < 1×10 □¹□). (G) GO Biological Process overrepresentation among differential peaks, ranked by GeneRatio. (H) Footprint (top) and occupancy (bottom) changes in acute (lower contrasts) and prolonged (upper contrasts) versus control. Middle bar: concordance (white), discordance (black) or non-significant (beige). Two-way ANOVA with Dunnett’s post-test (B, C) or two-tailed t test (D). (E–H) derive from two biological replicates per condition.

To discern whether these changes were accompanied by broader alterations in chromatin accessibility, we performed ATAC-seq on PMA/ionomycin restimulated cells exposed to either acute or prolonged lactate treatment. Across the 37,167 peaks detected, prolonged D-lactate altered the accessibility of only 315 peaks (0.84%) and acute D-lactate of 82 (0.22%), whereas neither prolonged nor acute L-lactate resulted in any differential peak with respect to control and with only 2 peaks differentially regulated between them (Figure 2E). Thus, despite a shared rapid increase in histone acetylation, measurable changes in chromatin accessibility were limited and restricted to D-lactate.

To further characterize the accessible regions altered by D-lactate, we performed HOMER motif enrichment analysis^39^, identifying 123, 9 and 135 differentially enriched motifs in prolonged D-lactate/control ((P)/Ctrl), acute D-lactate/control ((A)/Ctrl) and prolonged/acute D-lactate ((P)/(A)) contrasts, respectively. The (P)/Ctrl contrast was dominated by AP-1 and ETS modules, together with motifs linked to immune function (RUNX, STATs, NFAT and NF-κB) and mitochondrial metabolic/redox regulation (BACH1, GABPA, NRF2 and ESRRA). By contrast, (A)/Ctrl showed reduced accessibility at a subset of AP-1-associated motifs that became enriched after prolonged supplementation (Figure 2F). Overrepresentation analysis detected no significant terms among downregulated peaks, whereas upregulated peaks from both D-lactate conditions were associated with inflammatory response and immune activation; (P)/Ctrl was additionally enriched in apoptosis and ERK1/ERK2 cascade terms (Figure 2G). Thus, only D-lactate elicited detectable chromatin reorganization, which extended after prolonged exposure.

To obtain a broader view of transcription factor engagement, we applied TOBIAS across all 37,167 ATAC peaks, selecting transcription factors in the upper and lower tails of footprint change (q95 / q5) with high statistical support (q90)^40^. AP-1, ERK-associated and GC-rich-promoter-binding modules dominated the inferred regulatory landscape. Prolonged lactate exposure increased AP-1-related footprinting and occupancy, but with different kinetics: acute L-lactate initially reduced both modules, whereas acute D-lactate showed the opposite trend, which strengthened over time. Acute exposure also increased ERK-responsive and KLF/SP-family factors, although this response diminished after prolonged treatment, particularly with D-lactate.

Conversely, factors linked to type I interferon signaling (IRF and STAT families), cell identity (FOXC1 and FOXD3) and mitochondrial function (NRF1) showed early and sustained reductions (Figure 2H). Together, these findings indicate that lactate remodels transcription factor engagement through AP-1/ERK and type I interferon-related modules, combining shared and stereospecific features.

### D-lactate induces selective transcriptomic rewiring toward ERK/AP-1, metabolic, and redox programs

Given that chromatin accessibility changes were limited in extent but enriched in immune and AP-1/ERK-associated modules, we next interrogated how these alterations were integrated at the transcriptomic level. Cells were kept resting or restimulated with plate-bound anti-CD3 and anti-CD28 rather than PMA/ionomycin, to preserve early TCR-dependent signaling events that might otherwise be bypassed. We first inferred transcription factor activity from RNA-seq data through regulon analysis, capturing target-level regulatory changes. Overall, both lactate stereoisomers shared a substantial fraction of their regulatory output. Among the upregulated regulons were several immune-related transcription factors, including NFAT, NF-κB and ERK-responsive family members such as EGR1 and ETS1, irrespective of the activation state of the cells, as well as regulons linked to central metabolism control, including MYC, EPAS1 and HIF1A, whose direction and magnitude relative to control depended on the activation state and were more pronounced in D-lactate samples. In parallel, regulons associated with mitochondrial function and redox homeostasis, including GABPA, NFE2L2 and BACH1, were reduced. Type I interferon-associated regulons were consistently downregulated, reinforcing the pattern already suggested by the ATAC-based analyses, alongside a parallel decrease in proliferation-related modules (Figure 3A).

**Figure 3.**
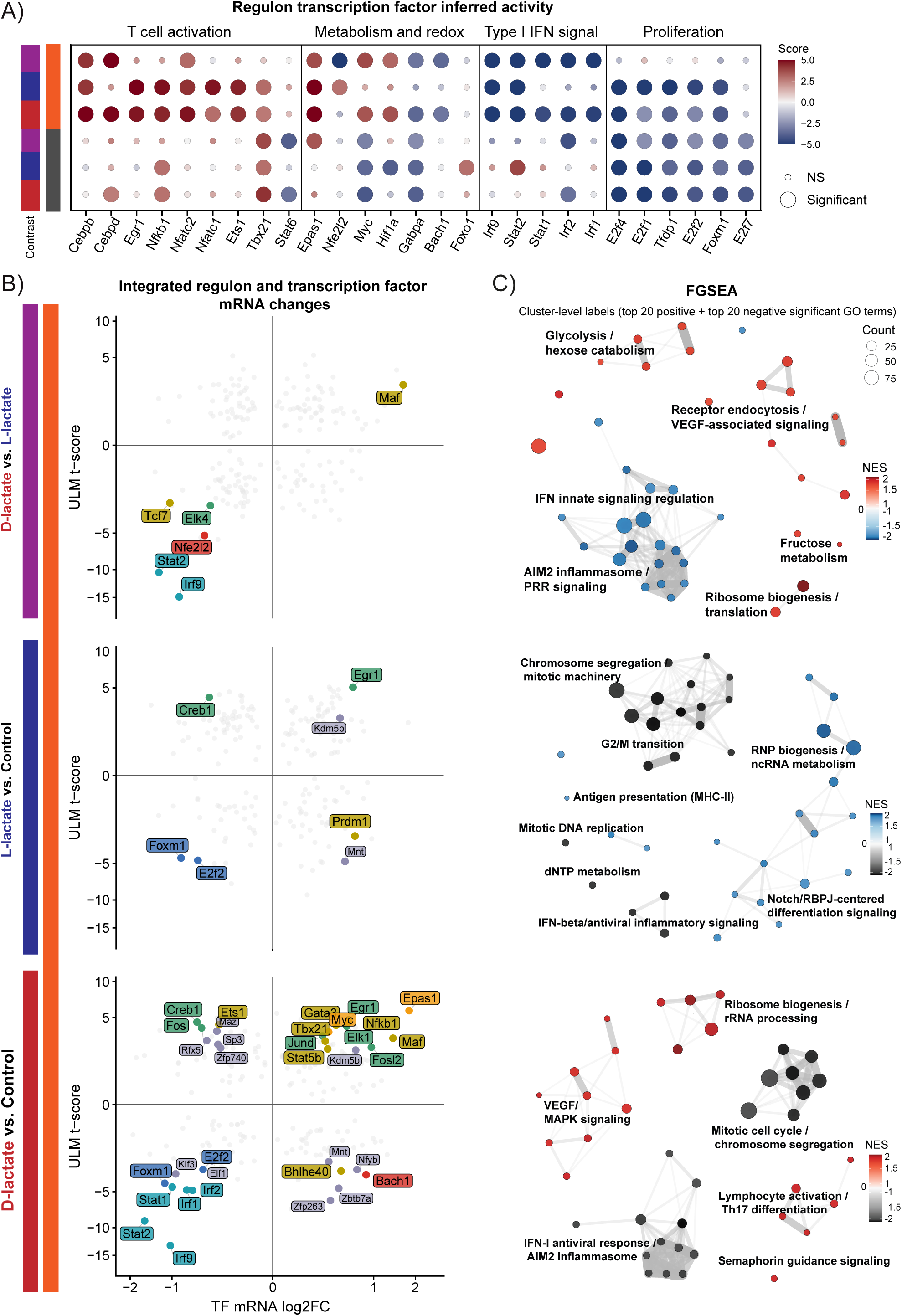
**Shared regulon responses are integrated into distinct process-level programs in lactate-exposed Th1 cells**. (A–C) Naïve WT CD4□ T cells were cultured for 5 days under Th1-polarizing conditions ± lactate, then stimulated with plate-bound αCD3 and αCD28 or kept resting, and processed for RNA-seq (two biological replicates per condition). (A) Bubble plot of the top 5 positively and negatively inferred regulons per contrast (ULM, DoRothEA), together with other differentially regulated immune-relevant regulons. (B) Dot plot of TFs reaching significance (p < 0.05) at both regulon-activity and mRNA expression levels in stimulated samples. (C) Clustered network of the top 20 positively and negatively enriched (NES) FGSEA GO terms in stimulated samples.

To refine this analysis, we integrated inferred transcription factor activity with changes in the transcript abundance of transcription factors, distinguishing concordant regulatory nodes from those likely driven by post-transcriptional or context-dependent mechanisms. Comparison of both layers revealed a shared regulatory signature, most clearly reflected in the downregulation of modules linked to cell cycle and type I IFN response. Beyond this common response, D-lactate displayed a broader spectrum of concordant changes relative to control than L-lactate, spanning metabolic regulators (MYC and EPAS1), redox-associated factors (BACH1 and NFE2L2), AP-1/ERK-associated factors (EGR1, JUN), and effector-linked pathways including NF-κB and NFAT (Figure 3B).

To resolve how these transcriptional changes were integrated into broader biological programs, we performed cluster-network analysis of the top 20 upregulated and downregulated FGSEA terms across the stimulated contrasts. This recapitulated the repression of proliferation and type I interferon-associated programs. Both stereoisomers also showed enriched ribosome-related programs but with distinct profiles: D-lactate was more strongly associated with rRNA maturation, ribosomal large subunit biogenesis and translation-related processes, whereas L-lactate was preferentially linked to ncRNA metabolism and ribonucleoprotein biogenesis. D-lactate further showed enrichment of multiple MAPK signaling and lymphocyte activation modules. Direct comparison between stereoisomers indicated that D-lactate was associated with a stronger repression of IFN-I and innate-related inflammatory response programs, together with a more pronounced reinforcement of ribosome-related clusters and clear enrichment of central carbon metabolism pathways, including sugar metabolism, in line with the preferential upregulation of MYC and EPAS1 at both the regulon and mRNA levels (Figure 3C). Thus, although both stereoisomers share a highly similar transcriptional regulatory network, this shared output is integrated differently at the pathway level, with the clearest differences involving sugar metabolism, ribosome/translation-related programs and type I IFN signaling.

### Lactate stereoisomers share ERK/AP-1-associated programs, whereas only D-lactate enhances ERK1/ERK2 phosphorylation

AP-1 and ERK-related modules were recurrently upregulated across ATAC-based footprinting and RNA-seq regulon data. In D-lactate samples this was particularly evident for ERK-related processes, including the GO term ‘ERK1 and ERK2 cascade’ (Figure 4A), although regulon and footprinting data indicated that ERK/AP-1-associated outputs were shared across both lactate conditions, albeit with different magnitude and configuration (Figure 4B). At the signaling level, phospho-ERK was selectively increased in D-lactate-treated cells relative to all other groups (Figure 4C). Because ERK1/ERK2 signaling regulates Perf-1 and GZMB expression in CD8^+^ cytotoxic T cells^41^, we tested its contribution to the D-lactate phenotype. Pharmacological inhibition with U0126 substantially impaired the ability of D-lactate to increase Perf-1 cell frequency and GZMB abundance, although it did not fully abrogate the effector burst, and modestly limited the ability of L-lactate to induce both molecules, consistent with a weaker ERK-dependent response (Figures 4D and 4E). We also considered Blimp-1 (*Prdm1*), which recruits T-bet to the *Gzmb* and *Prf1* loci^42^ and represses IFN-γ, as a candidate effector of the cytotoxic response. Although D-lactate induced Blimp-1, its deletion did not abolish the cytotoxic phenotype; instead, loss of Blimp-1 increased IFN-γ production under control and L-lactate conditions but not under D-lactate, consistent with reduced sensitivity to Blimp-1-mediated repression. Footprinting and expression data instead pointed to RUNX3 and EOMES as candidate effectors of the cytotoxic program (Figure S1).

**Figure 4.**
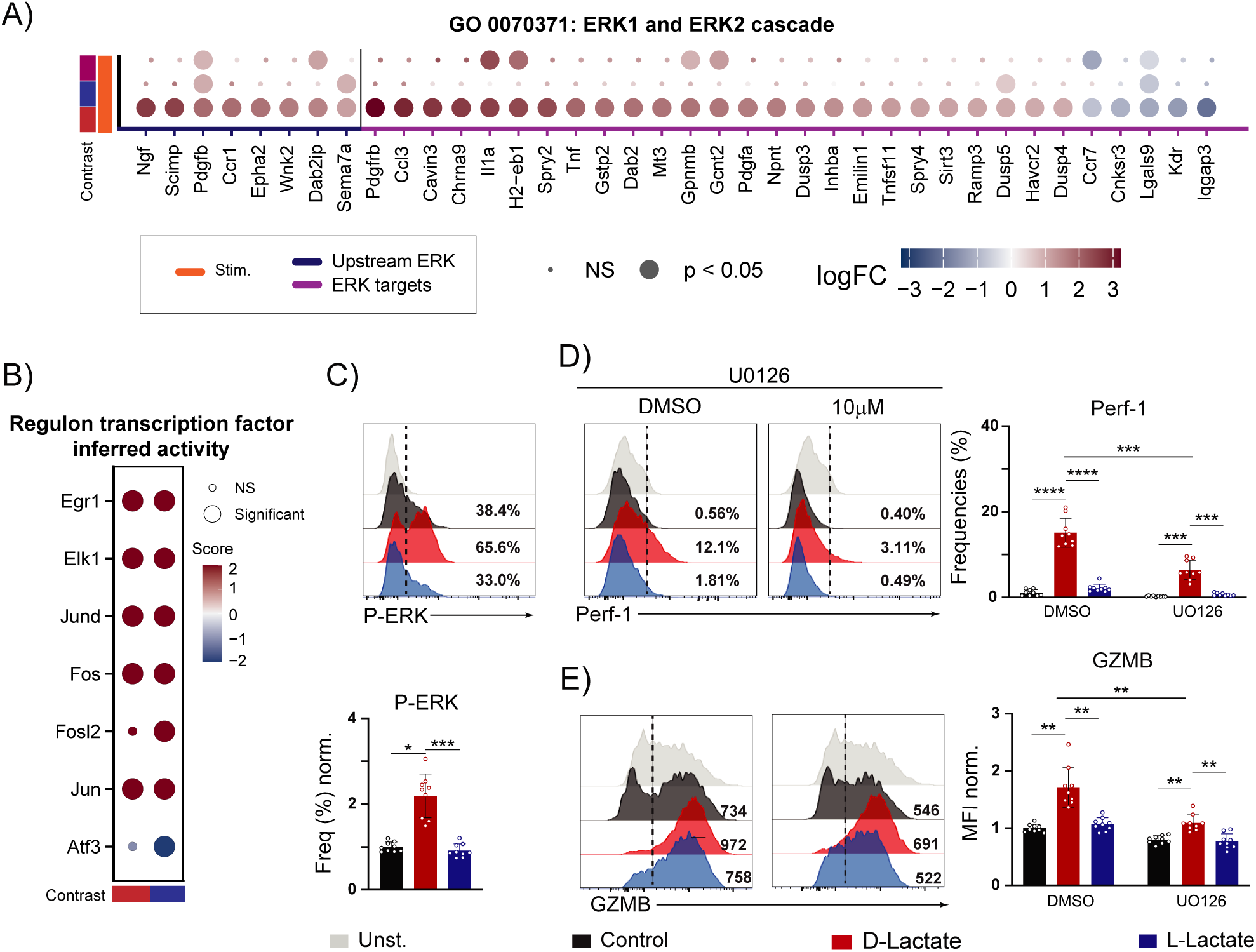
**ERK1/2 signaling contributes to lactate-driven cytotoxic programming and is preferentially enhanced by D-lactate in Th1 cells**. (A–E) Naïve WT CD4□ T cells were cultured for 5 (A–D) or 10 (E) days under Th1-polarizing conditions ± lactate. (D, E) DMSO or U0126 (10 μM) was added during restimulation. (A) limma-voom-normalized logFC values for differentially expressed genes (adjusted P < 0.05, |logFC| > 1) in GO:0070371, ERK1/ERK2 cascade. (B) ULM (DoRothEA) regulon analysis of significantly regulated AP-1 and ETS modules across contrasts (RNA-seq, two biological replicates per condition). (C) Representative histograms (top) and quantification (bottom) of P-ERK1/2 (n=9). (D) Representative histograms (left) and quantification (right) of Perf-1 (n=9). (E) Representative histograms (left) and quantification (right) of GZMB (n=9). One-way ANOVA with Tukey’s post-tests (C) or two-way RM-ANOVA with Tukey’s post-tests (D, E).

### D-lactate selectively enhances mTORC1 signaling and remodels central carbon metabolism in Th1 cells

Our transcriptomic data pointed to lactate-dependent changes in modules associated with anabolic signaling, ribosome biogenesis, glucose and carbon metabolism. Because mTORC1 is a central regulator of anabolic metabolism and promotes glycolysis and ribosomal protein synthesis through downstream targets such as S6^43^, we first assessed its activation: D-lactate-treated Th1 cells showed increased frequencies of phospho (P)-mTORC1 and P-S6 cells, while L-lactate had no observable effect (Figure 5A).

**Figure 5.**
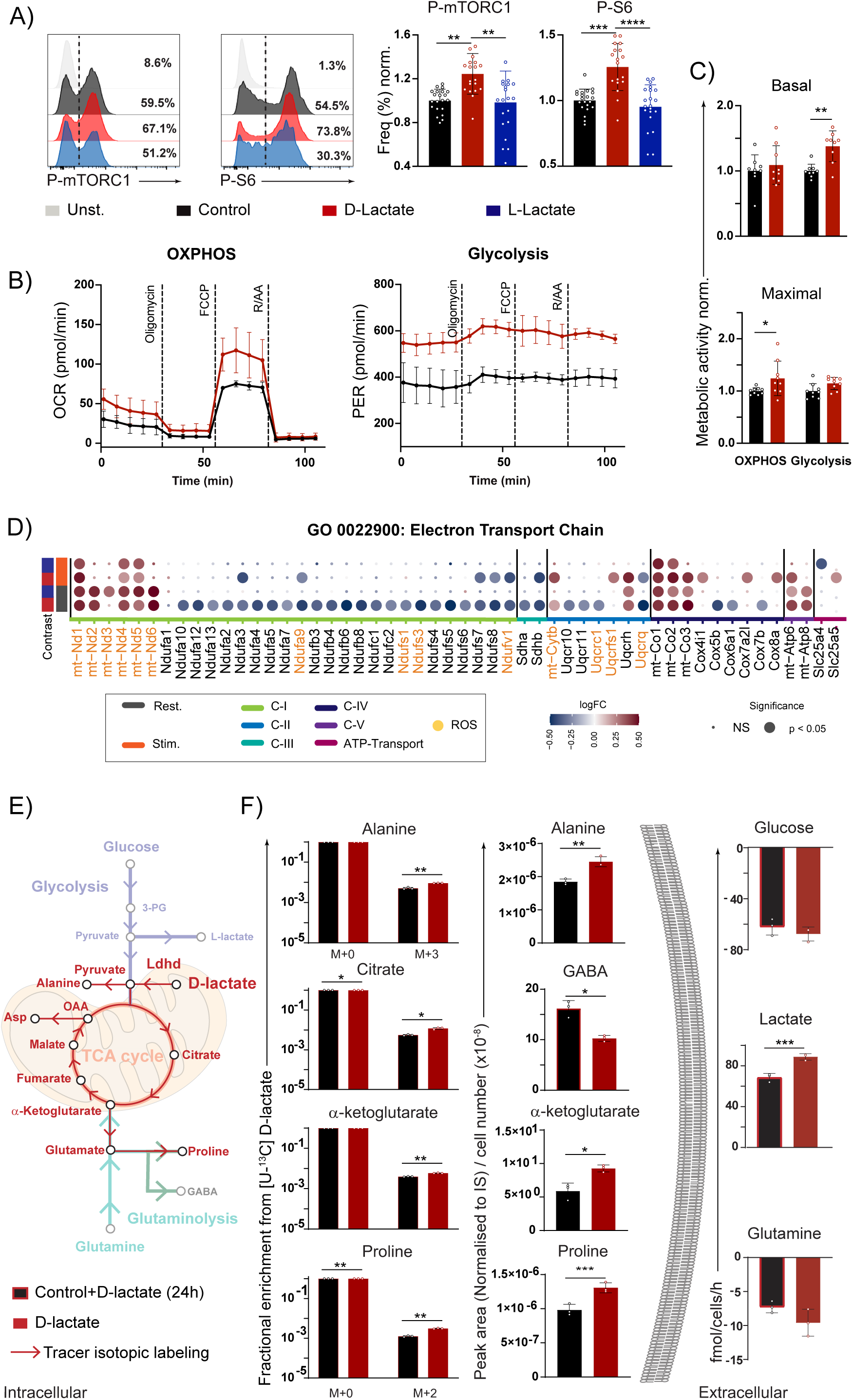
**D-lactate promotes metabolic reprogramming without serving as a major carbon source**. (A–D) Naïve WT CD4□ T cells were cultured for 5 days under Th1-polarizing conditions ± lactate. (E, F) On day 5, control and D-lactate-treated cells were replated in RPMI containing [U-¹³C]-D-lactate for 24 h. (A) Representative histograms (left) and quantification (right) of P-mTORC1 (S2448) and P-S6 (S240/244) (n=21). (B) Representative Seahorse kinetics of OXPHOS and glycolysis (n=3). (C) Basal (top) and maximal (bottom) activity in each pathway (n=9). (D) limma-voom-normalized expression values for differentially expressed genes (adjusted P < 0.05) within GO:0022900, electron transport chain (RNA-seq, two biological replicates per condition). (E) Schematic of central carbon metabolism. (F) Fractional ¹³C labeling (M+2 or M+3 isotopologues; left), intracellular pool abundance (center) and extracellular fluxes (right) of TCA-cycle intermediates and amino acids (n=3). RM-One-way ANOVA with Tukey’s (A), two-way ANOVA with Sidak (C, F (left)), or paired two-tailed t tests (F, center and right).

To determine whether mTORC1 contributed to the D-lactate-associated phenotype, we inhibited it with rapamycin during cytokine restimulation. Rapamycin did not eliminate the increase in IFN-γ frequencies between D-lactate and control samples, but completely abrogated the D-lactate-mediated increase in IFN-γ MFI (Figure S2A), and significantly reduced the upregulation of Perf-1 frequencies and GZMB MFI on day 10 (Figure S2B). mTORC1 therefore contributes to the functional effects induced by D-lactate but does not account for the full phenotype.

Because mTORC1 signaling promotes glycolytic and anabolic programs through MYC- and HIF-associated pathways ^44,45^, both of which were significantly upregulated in our regulon analyses under D-lactate conditions, we next assessed bioenergetic activity. Seahorse analysis revealed that D-lactate samples displayed an approximately 40% increase in basal glycolytic levels together with enhanced maximal respiratory capacity relative to control samples (Figures 5B and 5C).

The increase in maximal OXPHOS was notable, given the reduced NRF1 footprinting and decreased GABPA regulon activity identified above, two major regulators of nucleus-encoded respiratory genes^46,47^. However, the RNA-seq data revealed that lactate did not induce a uniform repression of the ETC but an asymmetric remodeling of ETC-subunit expression, with upregulation of multiple mitochondrion-encoded transcripts across several complexes together with downregulation of numerous nucleus-encoded subunits. This effect was most pronounced under D-lactate resting conditions and particularly affected Complex I, including ROS-relevant nuclear subunits such as *Ndufv1* and *Ndufs1* (Figure 5D).

To test whether D-lactate conferred a metabolic advantage over L-lactate during stimulation, as suggested by our FGSEA data, we tracked metabolic activity in cells stimulated in situ during Seahorse measurements. D-lactate-treated cells showed higher OXPHOS than their L-lactate counterparts, whereas glycolytic activity was very similar. Upon mitochondrial perturbation by oligomycin and R/AA, however, D-lactate samples exhibited greater glycolytic plasticity, consistent with enhanced mTORC1 activation and MYC regulon activity (Figure S2C). Acute removal of D-lactate during the assay caused a marked collapse in mitochondrial respiration and loss of this plasticity (Figure S2D). Although NRF1 footprinting decreased in both score and occupancy during lactate treatment, D-lactate was associated with enrichment of ESRRA- and ESRRG-linked signatures, two regulators connected to mitochondrial oxidative metabolism and respiratory gene programs^48,49^ (Figure S2E). Thus, during stimulation D-lactate reinforces mitochondrial oxidative metabolism while sustaining a glycolytic activity comparable to L-lactate, and increases the plasticity of the cells to upregulate glycolysis under metabolic stress.

Mammals express a stereospecific D-lactate dehydrogenase (LDHD) that converts D-lactate into pyruvate, potentially allowing its use as a precursor for macromolecule biosynthesis ^50,51^. Although *Ldhd* expression levels were modest under our experimental conditions, *Ldhd* mRNA levels were increased approximately sevenfold in D-lactate-treated samples (Figure S2F). We therefore investigated whether D-lactate was metabolized in Th1 cells by performing tracing experiments in cells previously cultured under control or D-lactate conditions and then exposed to [U-¹³C] D-lactate for 24 hours.

We detected low but consistent incorporation of D-lactate-derived carbon into TCA-cycle intermediates (Figures 5E and 5F), representing approximately 0.2–1.2% of total metabolite pools. Labelling was higher in D-lactate-differentiated cells, with significantly increased citrate (M+2) and α-ketoglutarate (M+2), but not downstream metabolites. D-lactate-derived carbon was also detected in proline (M+2) and alanine (M+3). The increase in alanine M+3 was consistent with increased conversion of D-lactate into pyruvate and subsequent transamination to alanine, providing indirect evidence of increased *Ldhd* activity. Thus, CD4□ T cells metabolize D-lactate, but its minimal anaplerotic contribution is unlikely to explain the hyperfunctional phenotype.

After the 24-hour pulse, D-lactate-treated cells retained metabolic differences from control, indicating that short-term treatment did not reproduce the state induced by prolonged exposure. α-Ketoglutarate was the only TCA intermediate with an increased pool. With glutamine consumption unchanged, the decrease in GABA alongside increased proline likely reflects redirection of glutamate-derived carbon toward proline at the expense of GABA. Extracellular (unlabeled) lactate was also elevated, consistent with increased glycolytic output. Functionally, 24-hour D-lactate-pulsed cells showed reduced GZMB and IFN-γ production in strong contrast to cells differentiated with D-lactate (Figure S2G). Thus, prolonged D-lactate exposure therefore rewires central carbon metabolism rather than serving primarily as an oxidative substrate (Figures 5F and S2G).

To determine whether the augmented metabolism induced by D-lactate contributed to its effects on cytokine production, we inhibited glycolysis or OXPHOS with 2-deoxyglucose (2-DG) or rotenone/antimycin A (R/AA), respectively. Inhibition of either pathway did not eliminate the differences in IFN-γ and GZMB expression between D-lactate-treated and control samples but instead increased the disparity between them (Figure S3A), indicating that the phenotype cannot be explained solely by differences in glycolytic or oxidative metabolism and raising the possibility that lactate confers an advantage under metabolically restrictive conditions. To test this, we reduced glucose concentration in the culture medium to 1 mM. Because limited cell recovery precluded Seahorse analysis, we used mTORC1 signaling and mitochondrial parameters as proxies for metabolic fitness. Phosphorylation of mTORC1 and S6, together with P-ERK1/ERK2, remained selectively enhanced in D-lactate-treated cells and became even more pronounced under glucose restriction, as did mitochondrial TMRM/MitoSpy ratios (Figures S3B–S3E). Accordingly, after 3 days of glucose restriction, D-lactate still sustained enhanced expression of IFN-γ, TNF-α, GZMB and Perf-1, whereas L-lactate produced only a modest increase in GZMB. This effect was lost when cells were deprived of their respective lactate, arguing against the establishment of a stable cell-intrinsic memory of lactate-induced phenotypic changes (Figures S3F and S3G). Taken together, under metabolically restrictive conditions D-lactate sustains key proinflammatory and cytotoxic molecules while preserving mTORC1, ERK1/ERK2 signaling and mitochondrial fitness.

### D-lactate redirects redox responses and depends on glyoxalase to limit mitochondrial ROS

Regulon inference suggested reduced activity of NFE2L2 and BACH1, two central regulators of antioxidant and heme/redox stress responses, respectively^52,53^. However, motif enrichment analysis of differential peaks together with TOBIAS footprinting pointed to increased NFE2L2-associated chromatin engagement (Figures 2F, 2H and S4A), arguing against a uniform activation or repression of this pathway. To dissect the redox response in detail, we generated a heatmap based on three ontology/pathway categories containing NFE2L2-associated targets: the Reactome KEAP1–NFE2L2 pathway, glutathione metabolism, and oxidative stress response.

This revealed that the NFE2L2-associated response was not uniformly activated or repressed in D-lactate-treated cells, but selectively redistributed across downstream branches. *Nfe2l2* itself was consistently downregulated across all D-lactate comparisons, and several canonical effector branches were attenuated, most notably the proteostatic and glutathione-producing modules. Among the genes that remained induced were *Hmox1* together with cytosolic electrophile-detoxification/GSH-consuming genes, including multiple members of the *Gstm* family, as well as extracellular GSH-recycling genes such as *Ggt1* and *Ggt5*. Conversely, genes involved in the detoxification of cytoplasmic and mitochondrial ROS, exemplified by *Cat* and *Sod2*, respectively, were downregulated. By contrast, L-lactate-treated samples did not display a comparably coherent pattern of oxidative stress gene regulation (Figure 6A). These results argue against a canonical NFE2L2-driven antioxidant program and support a branch-specific redirection of the redox response.

**Figure 6.**
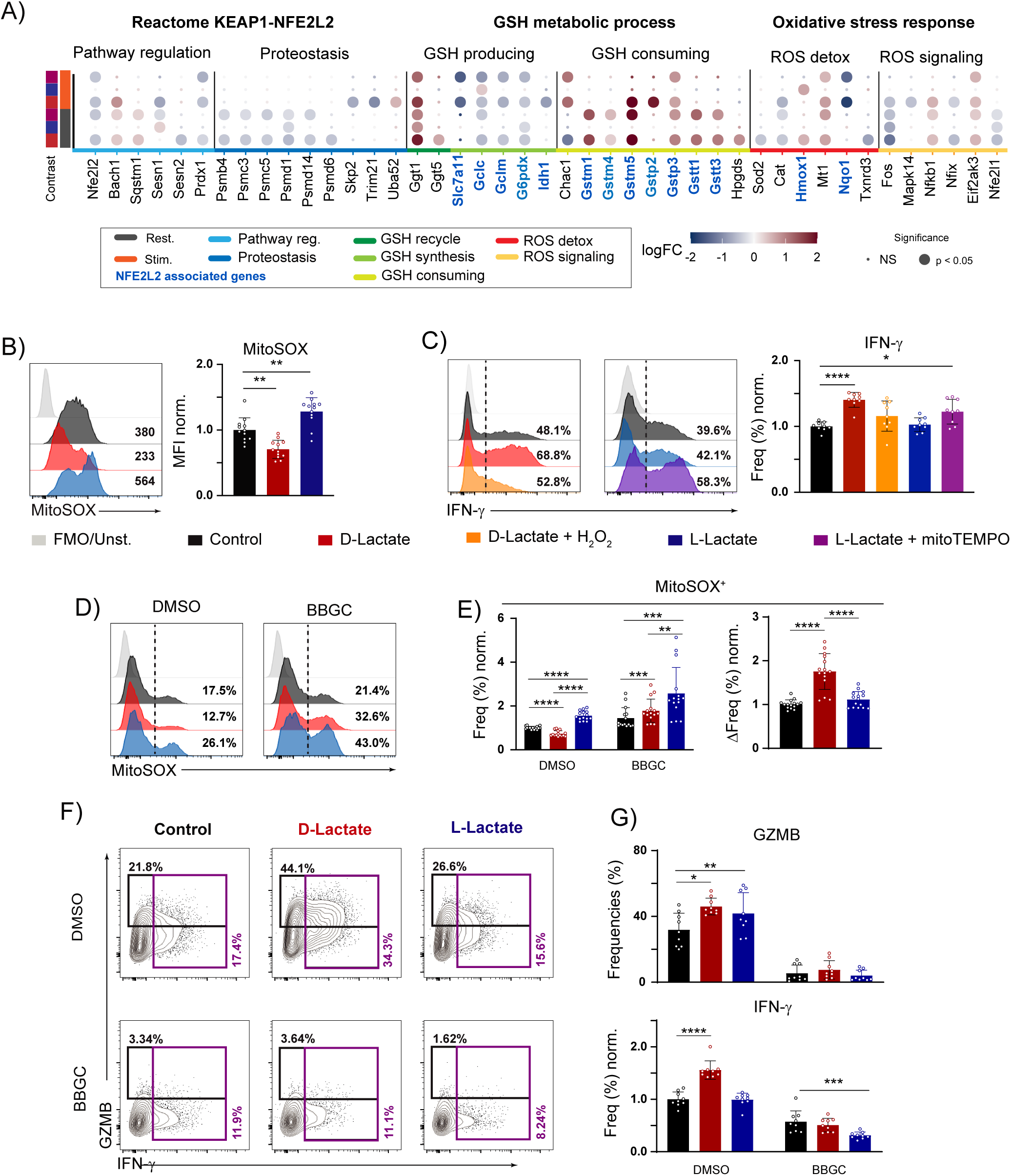
**D-lactate depends on the glyoxalase system to preserve low mitochondrial ROS and sustain effector function**. (A–G) Naïve WT CD4□ T cells were cultured for 5 days under Th1-polarizing conditions ± lactate. (A) limma-voom-normalized logFC values for differentially expressed genes (adjusted P < 0.05, |logFC| > 0.4) in the Reactome KEAP1-NFE2L2 pathway, GO glutathione metabolic process and WikiPathways oxidative stress response gene sets (RNA-seq, two biological replicates per condition). (B) Representative histograms (left) and MitoSOX MFI (right) (n=12). (C) Representative histograms (left) and quantification (right) of IFN-γ; mitoTEMPO (2 μM) or H O (10 μM) added during PMA/ionomycin restimulation (n=9). (D) Representative MitoSOX histograms after BBGC (1 μM, 30 min). (E) Quantification of (D) (n=15). (F) Representative contour plots of IFN-γ and GZMB after BBGC (1 μM, 4 h). (G) Quantification of (F) (n=15). One-way ANOVA with Dunnett’s post-tests (B, C) or two-way ANOVA with Tukey’s post-tests (E, G).

This was particularly relevant given the altered mitochondrial:nuclear transcriptional stoichiometry, especially in Complex I, where several subunits directly implicated in ROS generation were stereospecifically downregulated upon D-lactate treatment. We therefore assessed the generation of different mitoROS species. D-lactate limited mitochondrial O□□ formation relative to control samples, whereas L-lactate significantly increased it (Figure 6B). To determine whether the reduction in O□□ could reflect increased conversion into H□ O□, we quantified mitochondrial H□ O□ production and observed the same overall trend (Figure S4B). Thus, lactate stereoisomers exert opposite effects on the generation of ETC-derived mitoROS.

We next asked whether these differential mitoROS patterns contributed to their resulting phenotypes, experimentally increasing or scavenging ROS levels in each condition with H□ O□ or the mitochondria-targeted scavenger mitoTEMPO, respectively. When H□ O□ was added to D-lactate-treated cells, the frequency of IFN-γ cells was reduced toward control levels; conversely, mitoROS scavenging in L-lactate-treated samples increased IFN-γ levels compared to untreated L-lactate cells (Figure 6C).

We had also observed that acute D-lactate treatment of control cells reduced both GZMB and IFN-γ production, mirroring the effect of SAHA. MitoSOX staining revealed that acute exposure to either lactate stereoisomer increased superoxide production, which was reverted by mitoTEMPO; accordingly, IFN-γ was reduced under the same conditions and rescued by mitoTEMPO scavenging (Figure S4C). Consistent with this effect, SAHA also increased mitochondrial superoxide (Figure S4D). Together, these data link mitoROS levels to the capacity of lactate stereoisomers to modulate cytokine production.

D-lactate increased *Ldhd* expression despite the minimal incorporation of labeled D-lactate into downstream metabolites which, together with the enhanced glycolytic activity of these cells, suggested that *Ldhd* upregulation might reflect a broader metabolic adaptation rather than the use of D-lactate as a bioenergetic substrate. Because D-lactate is the end product of the glyoxalase pathway, composed of Glo1 and Glo2, which detoxifies the glycolytic byproduct MG, we next asked whether this system contributed to the D-lactate phenotype. Intracellular flow cytometry revealed no significant changes in Glo1 expression but a significant increase in Glo2 levels in D-lactate-treated samples (Figures S4E and S4F). MG-Lys levels were lower in D-lactate-treated cells under steady-state conditions and after 24 hours of exogenous MG exposure, consistent with more efficient handling of dicarbonyl stress (Figure S4G). Because the glyoxalase system has been linked to mitochondrial redox regulation beyond MG detoxification, including modulation of the glutathione pool and protein glutathionylation^54–56^, we inhibited Glo1 with BBGC. Glyoxalase inhibition caused D-lactate-treated cells to lose their ability to limit mitochondrial ROS and led to the strongest increase in mitochondrial superoxide among all conditions (Figures 6D and 6E), abrogating the D-lactate-mediated effect on IFN-γ and GZMB production (Figures 6F and 6G). Hence, D-lactate-treated cells are particularly dependent on this axis to preserve mitochondrial redox balance and sustain effector function.

### D-lactate reinforces proinflammatory, anabolic and histone acetylation programs in human CD4^+^ T cells

To extend our findings on the stereospecific regulatory effects of D-lactate from murine to human cells, we sorted naïve CD4□ T cells from healthy individuals and polarized them towards Th1 *in vitro*. Consistent with our murine findings, D-lactate increased the frequencies of IFN-γ- and TNF-α-producing cells, and a significant increase in MFI was observed for IFN-γ, while L-lactate had no observable effect (Figures 7A and 7B). Moreover, D-lactate selectively increased histone H3 acetylation at K9 and K27 (Figures 7C and 7D). At the metabolic level, D-lactate-treated cells showed increased glycolysis and OXPHOS relative to control, although the OXPHOS difference did not reach statistical significance. As in murine cells, removal of D-lactate was associated with lower OXPHOS, while glycolysis remained largely unaffected; L-lactate caused only minor, non-significant increases in both pathways (Figures 7E and 7F). Lastly, both lactates increased TMRM frequencies, whereas only D-lactate increased the P-mTORC1/mTOR ratio (Figures 7G-7I). Together, these data indicate that the proinflammatory and anabolic response to D-lactate, along with its effect on histone acetylation, is conserved in human CD4□ T cells.

**Figure 7.**
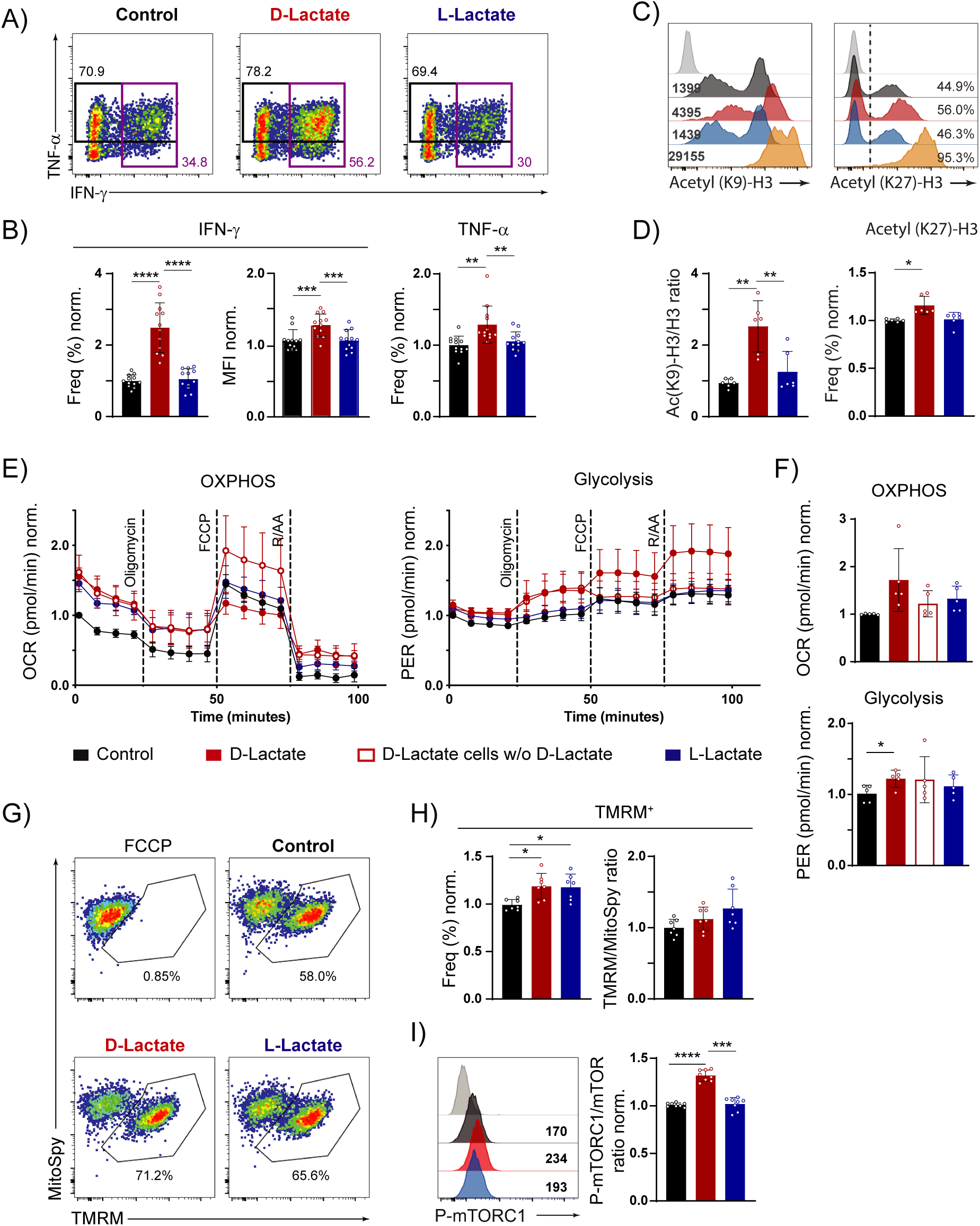
**D-lactate reproduces the proinflammatory and anabolic response in human CD4□ Th1 cells**. (A–I) Naïve human CD4□ T cells were cultured for 5 days under Th1-polarizing conditions ± lactate. D-lactate was withdrawn from a fraction of D-lactate-differentiated cells during Seahorse measurements. (A) Representative contour plots of IFN-γ and TNF-α. (B) Quantification of (A) (n=12). (C) Representative histograms of Acetyl(K9)-H3 and Acetyl(K27)-H3. (D) Quantification of (C) (n=6). (E) Representative Seahorse kinetics of OXPHOS and glycolysis (n=5). (F) Basal OXPHOS (top) and glycolytic (bottom) activity (n=5). (G) Representative pseudocolor plot of TMRM and MitoSpy stainings. (H) Quantification of (G) (n=7). (I) Representative histogram (left) of P-mTORC1 and P-mTORC1/mTOR ratio (right) (n=8). Data are mean ± SD, except (E) ± SEM. RM-One-way ANOVA with Tukey’s post-tests.

## Discussion

By directly comparing lactate stereoisomers during Th1 cell differentiation, we found that they do not simply produce the same response to different degrees but instead drive qualitatively distinct effector states. Both promoted the acquisition of cytotoxic features, but D-lactate did so much more strongly, while L-lactate had only a modest effect. The difference was even more evident in the inflammatory arm of the response: D-lactate increased TNF-α and IFN-γ production in both in murine or human Th1 cells. In contrast, L-lactate either reduced the production of both cytokines or left it unaffected in murine or human Th1 cells, respectively.

As expected, rapid histone hyperacetylation was an early response shared by both stereoisomers. However, it was insufficient to promote cytokine production, as SAHA failed to reproduce the phenotype and *Ifng* transcript levels increased only after prolonged D-lactate exposure. In parallel, only D-lactate altered chromatin accessibility, and although the changes were modest in magnitude, acute and prolonged differential peaks were enriched near genes associated with immune activation and inflammatory responses. Motif enrichment, GOSEA and footprinting analyses converged on AP-1/ERK-associated modules. Consistent with these observations, D-lactate stereospecifically activated ERK1/ERK2 signaling, which regulates AP-1 and ETS-associated factors including JUN and EGR family members^57^. Pharmacological ERK1/ERK2 inhibition impaired acquisition of cytotoxic features in D-lactate-treated cells, supporting a functional contribution of this pathway.

In parallel, the persistent repression of STAT/IRF-associated programs across chromatin, regulon, and pathway-level analyses indicated that D-lactate does not simply amplify canonical Th1 cell activation. Consistent with this, its ability to enhance IFN-γ production not only persisted but became even more pronounced in *Stat1*-, *Stat4*- and *Tbx21*-deficient cells, arguing that the D-lactate-associated inflammatory phenotype is not primarily mediated through canonical STAT1/STAT4/T-bet regulation.

The acute D-lactate condition provided an important indication of mechanistic hierarchy. Although short-term exposure was sufficient to induce histone hyperacetylation and to initiate selective chromatin remodeling toward proinflammatory/AP-1/ERK-associated programs, it did not promote the effector state and instead reduced IFN-γ and GZMB expression. In contrast, direct modulation of mitochondrial ROS under the same conditions was sufficient to rescue or impose cytokine production, indicating that mitochondrial ROS control is more proximally linked to effector function than chromatin permissiveness.

No single master regulator of the cytotoxic CD4□ T cell program has been identified, but several transcription factors have been linked to the acquisition of cytotoxic features. Blimp-1 was a plausible candidate, but its ablation did not abolish the increase in cytotoxic molecules induced by D-lactate. Its clearest effect was on the inflammatory arm: loss of Blimp-1 increased IFN-γ production under control and L-lactate conditions but had no effect in D-lactate-treated cells, indicating that D-lactate already relieves Blimp-1-mediated repression, an interpretation supported by the reduced Blimp-1 footprinting and occupancy observed despite higher Blimp-1 expression. RUNX3 and EOMES, previously implicated in the acquisition of cytotoxic features by Th1-derived cytotoxic CD4□ T cells^34^, are more likely consolidators of the program: lactate increased both their protein levels and their footprinting signals, most strongly under D-lactate, which also broadened their transcriptomic output, whereas L-lactate reduced occupancy at their target sites.

D-lactate also induced an anabolic program evident in increased mTORC1 and S6 phosphorylation, enrichment of ribosome biogenesis and translation-related pathways, and stronger *Myc* and *Epas1* regulon activity. These changes align with higher basal glycolysis and metabolic plasticity. Still, this reinforcement was insufficient to explain the phenotype, since glycolysis or OXPHOS inhibition did not eliminate the D-lactate advantage and rapamycin only partially reduced IFN-γ, GZMB and Perf-1. Thus, this anabolic state supports but does not determine the hyperfunctional phenotype.

Beyond this anabolic program, the stereospecific D-lactate phenotype could not be explained by its contribution as an anaplerotic source of carbon or a bioenergetic role. Isotopic tracing experiments showed only minimal contribution of D-lactate-derived carbon to TCA-related metabolites, and this incorporation was higher in D-lactate-differentiated cells, consistent with their elevated *Ldhd* expression. The carbon contribution of D-lactate metabolism is therefore far too small to sustain the observed phenotype and *Ldhd* upregulation reflects a broader metabolic adaptation rather than the exploitation of D-lactate as a fuel. More than metabolic activity per se, the key feature underlying the stereospecific phenotypic outcomes was the distinct regulation of mitochondrial ROS by each lactate enantiomer. Consistent with earlier studies showing that L-lactate can promote mitoROS generation as a consequence of its own metabolism^58^, D-lactate instead limited mitoROS under our experimental conditions, and modulation of mitoROS produced during restimulation was sufficient to revert or impose the proinflammatory phenotype in D- and L-lactate, respectively.

Our data indicate that lactate supplementation reshapes mitochondrial biology, as reflected by ETC transcriptional stoichiometric remodeling consistent with reduced NRF1 footprinting and GABPA regulon activity. Both enantiomers increased transcription of mitochondrially encoded ETC subunits across complexes I, III, IV and V, but only D-lactate consistently downregulated nucleus-encoded complex I subunits. This mitonuclear remodeling resembles the process described in long-lived mice, where lower abundance of nucleus-encoded matrix-arm complex I subunits is associated with improved assembly and reduced superoxide production^59,60^. Because NDUFV1 and NDUFS1 reside in the matrix arm, a major site of superoxide generation, such remodeling could constrain ROS generation without impairing electron transport, consistent with reports that D-lactate can limit mitoROS under suboptimal complex I activity^61^. Although individual ETC complex activities were not measured, Seahorse and mitochondrial staining showed preserved mitochondrial fitness: D-lactate-treated cells displayed increased maximal OXPHOS and membrane potential, alongside reduced mitochondrial superoxide and H O . Notably, this reduction occurred without a uniform NFE2L2 signature. The lack of a reinforced glutathione-based antioxidant program argues against enhanced ROS buffering as the cause of the lower ROS levels, pointing instead to reduced mitochondrial ROS production at the ETC as the primary mechanism.

The glyoxalase system plays a dual role in mitigating metabolic stress, being central to the regulation of mitochondrial redox balance and also bridging glycation stress through its critical role in MG detoxification^56^. D-lactate increased basal glycolytic rates by more than 40% while reducing MG-derived protein adduct formation, despite the increased flux. Because MG accumulation impairs the activity of complexes I and III, promoting mitochondrial superoxide generation^62,63^, its effective neutralization protects the ETC from a second, glycation-derived source of oxidative stress. Accordingly, MG accumulation upon glyoxalase inhibition increased mitochondrial superoxide production, reaching its maximum in D-lactate-treated cells in line with their higher glycolytic flux. This inhibition also abrogated the D-lactate-driven increase in IFN-γ and GZMB. Enhanced glyoxalase activity is therefore required for the low mitoROS state that sustains the D-lactate phenotype.

Glucose restriction in chronic inflammation and tumors impairs T cell effector function and promotes exhaustion^64–67^. Notably, D-lactate sustained cytokine production, mitochondrial membrane potential, and mTORC1/ERK signaling under glucose-restricted conditions. Moreover, targeted D-lactate delivery has reprogrammed M2 toward M1 macrophages and enhanced immunotherapy in hepatocellular carcinoma^32^. Redirecting such platforms toward CD4□ T cells might complement macrophage reprogramming by reinforcing local inflammatory and cytotoxic responses. These findings may thus extend to settings in which D-lactate does not arise naturally, pointing to a potential therapeutic use.

Some limitations of our study should be acknowledged. The data were generated primarily in murine Th1 cells spanning different murine strains and stimulation settings. Although our human CD4□ T cell experiments reproduced the proinflammatory and anabolic arms of the response, whether the cytotoxic and redox components extend to human cells remains to be determined. In addition, although our results indicate that mitochondrial ROS are functionally epistatic to histone hyperacetylation and early chromatin rewiring in the acquisition of the effector phenotype, the precise integration of this redox layer with downstream signaling pathways and transcription factor engagement remains to be fully resolved.

Overall, our findings show that D- and L-lactate are not functionally interchangeable during Th1 cell differentiation, but instead bias cells toward clearly different effector states. Although D-lactate drives higher glycolytic and mitochondrial metabolic rates, convergent metabolic adaptations result in a lower oxidative burden. ETC-subunit transcriptional remodeling was associated with reduced mitoROS, while glyoxalase-dependent buffering limited the mitochondrial stress associated with elevated glycolytic activity. By establishing this redox state, D-lactate reinforces both inflammatory and cytotoxic functions and helps to sustain effector activity under metabolically restrictive conditions such as glucose scarcity. Our results thus expand current understanding of the mechanisms that favor acquisition of cytotoxic CD4□ T cell features and provide a framework for how metabolite stereochemistry shapes effector differentiation, indicating that metabolic adaptation in T cells is not simply a matter of increasing bioenergetic output, but also of how redox balance and stress resilience are maintained.

## Materials and methods

### Mice

C57BL/6J or C57BL/6N mice were used as wild type (WT) controls. In addition, strains carrying the following genetic modifications were used: *Stat1*^-/-^, *Stat4*^-/-^, *Tbx21*^-/-^, Blimp-1eGFP, SMARTA^+^ or *Il7raCrexPrdm-1*^fl/fl^ (Blimp-1^fl/fl^xCre from now on) and *Tcrbd*^-/-^ mice. Blimp-1eGFP and Blimp-1^fl/fl^xCre mice were obtained from Prof. Thomas Dörner and Prof. Andreas Diefenbach, respectively. SMARTA^+^ and Blimp-1^fl/fl^xCre mice were bred in a heterozygous state for their respective transgene (SMARTA and *Il7ra*Cre, respectively). For all the experiments, male and female mice were used at an age of 8 to 12 weeks. All animals were bred under specific-pathogen free (SPF) conditions in the animal facility of Charitè – Universitätsmedizin Berlin.

### Human samples

Human buffy coats from healthy donors were obtained from the German Red Cross (DRK), and donors provided written informed consent. Their use was approved by the ethics committee of Charitè – Universitätsmedizin Berlin.

### T cell isolation and cell sorting

Mouse and human CD4^+^ T cells were isolated from spleens or buffy coats, respectively. Samples were enriched for CD4□ using CD4-biotin (L3T4, BD) or hCD4□ microbeads (Miltenyi) and subsequently stained with the following antibody mixes: CD8a (53-6.7, BD), CD25 (pC61.5, DRFZ), CXCR3 (CXCR3-173, eBiosciences), CD62L (MEL-14, eBiosciences), CD44 (IM7, DRFZ) or hCD4□ (OKT4, BioLegend), hCD25 (BC96, eBiosciences), hCD45RA (4G11, DRFZ) / hCD127 (A019D5, BioLegend) hCD45RO (UCHL1, DRFZ), hCXCR3 (G025H7, BioLegend), hCRTH2 (BM16, Miltenyi). We defined naïve mouse and human CD4^+^ T cells as: (CD4^+^ propidium iodide^-^ CD25^-^ CXCR3^-^CD62L^+^CD44^-^) or (CD4^+^ propidium iodide^-^ CD25^low^CXCR3^-^CRTH2^-^ CD45RO^-^ and either CD45RA^+^ or CD127^+^), respectively.

### Activation and differentiation of CD4^+^ T cells with antibodies or antigenic peptide

Mouse and human CD4^+^ T cells were stimulated for 48h with plate-bound anti-CD3 (145-2C11, eBiosciences) and anti-CD28 (CD28.6, eBiosciences), or h-αCD3 (UCHT1, eBiosciences) and h-αCD28 (CD28.2, eBiosciences). SMARTA^+^ T cells were alternatively stimulated with irradiated splenocytes from TCRβδ-deficient mice loaded with GP61 peptide (GLKGPDIYHGVYQFKSVEFD, 1μM; R.Volkmer, Charitè) at a 4:1 ratio (APC : T cell). All samples were cultured with RPMI 1640+GlutaMax-I (Gibco) containing: 10% (v/v) FCS (Gibco), penicillin (100 U/ml; Gibco), streptomycin (100 μg/ml; Gibco), ß-mercaptoethanol (50 μM; Sigma). For murine Th1 differentiation, cultures were supplemented with IL-2 (5ng/mL; Miltenyi), IL-12 (5ng/mL; Miltenyi) and anti-IL-4 (11b11; 10μg/mL; DRFZ). For human Th1 differentiation cultures were supplemented with hIL-12 (5ng/mL), h-anti-IL-4 (10μg/mL) and hIL-2 (5ng/mL). Some samples were additionally supplemented with either Na-D-lactate (20mM) or Na-L-lactate (20mM; both from Sigma-Aldrich). Most of the analyses were carried out either after one (5 days) or two rounds (10 days) of *in vitro* stimulation. For some experiments, MG (500μM; Sigma-Aldrich) was added for 24h from day 4 to 5. In glucose-restricted experiments, day 5 differentiated cells were resuspended in Agilent SeaHorse XF RPMI medium supplemented with 10% (v/v) FCS (Gibco), penicillin (100 U/ml; Gibco), streptomycin (100 μg/ml; Gibco), ß-mercaptoethanol (50 μM; Sigma), glutamine (Gibco, 2mM), Na-pyruvate (Sigma, 1mM), glucose (Sigma, 1mM) and IL-2 (5ng/mL) for three days, during which medium was replenished daily.

### Restimulation and detection of intracellular cytokines and cytotoxic molecules

Intracellular detection of cytokines was performed by stimulating cells in RPMI medium containing PMA (5ng/mL) and ionomycin (500ng/mL, both from Sigma-Aldrich) for 4 hours. After the initial 30 minutes, brefeldin A (5μg/mL, Sigma-Aldrich) was added to the cultures until the end of the restimulation. For some of the experiments, cells were pre-incubated with mitoTEMPO (2μM) prior to stimulation or re-stimulated in the presence of H_2_O_2_ (10μM), rapamycin (100nM; Thermo-Fisher), U0126 (10μM; Bio Techne), SAHA (10μM; Bio Techne), BBGC (1μM; Sigma-Aldrich) for the entire re-stimulation time. Samples were stained with combinations of the following antibodies: CD4□ (L3T4/GK1.5, BD), IL-2 (JES-5H4, eBiosciences), TNF-α (MP6-XT22, Biolegend), GzmB (QA16A02, Biolegend), Perforin-1 (S1600, Biolegend), IFN-γ (XMG1.2, eBiosciences), hCD4□ (OKT4, BioLegend), hIFN-γ (B27, BD) and hTNF-α (Mab11, BioLegend).

### Transcription factor staining

Cells were fixed and permeabilized with the Foxp3/ Transcription Factor Staining Buffer Set (eBioscience) according to manufacturer’s protocol. Samples were stained in Permeabilization buffer 1X containing the following antibodies: CD4□ (RM4-5, eBiosciences) and Eomes (Dan11mag, eBiosciences).

### Intracellular staining of methanol-permeabilized samples

For intracellular detection of markers in methanol-permeabilized samples, cells were kept for 4 hours in RPMI medium alone or containing PMA (5ng/mL) and ionomycin (500ng/mL, both from Sigma). After that period, cells were fixed in 4% formaldehyde. Permeabilization was accomplished by adding 100% of ice-cold methanol to the fixed pellet. Samples were stained with different combinations of the following antibodies: Histone H3 (D1H2, CellSignaling), Acetyl-Histone H3 (Lys27) (D5E4, CellSignaling), Acetyl-Histone H3 (Lys9) (C5B11, CellSignaling), Phospho-mTOR(Ser2448) (MRRBY, Thermo-Fisher), Phospho-S6 (Ser240/244) (D68F8, Cell Signaling), Phospho-ERK (p44,p42) (Thr202/Tyr204) (197G2, CellSignaling), Runx3 (D6E2, Cell Signaling), Glo1 (6F10, abcam), Glo2 (ab154108, abcam) and CD4□ (RM4-5, eBiosciences) or hCD4□ (OKT4, BioLegend). For some of the markers a secondary staining was performed containing donkey anti-mouse IgG (AB_2556618; Invitrogen) and donkey anti-rabbit IgG (Polyclonal; Invitrogen) prior to acquisition.

### Mitochondrial stainings

Cells were resuspended in incomplete RPMI medium (lacking FCS) containing the various mitochondrial dyes. Samples were co-stained with MitoSpy Green FM (1μM, BioLegend) and TMRM (30nM, Thermo-Fisher). For O_2_ ^-^ and H O detection, cells were stained with MitoSOX (3μM, Thermo-Fisher) or MitoPY1 (5μM, Tocris), respectively.

### *In vitro* cytotoxic assay

SMARTA^+^ Th1 cells were differentiated for 5 days with or without lactate (D- or L-lactate, 20mM) as previously described in this section. On day 3, B cells were isolated from TCRβδ-deficient mice using the same methodology as for the CD4□ enrichment, exchanging only the primary antibody used for the positive magnetic selection to CD19 (ID3, BD). B cells were then plated in RPMI supplemented with dextran sulfate (25μg/mL, Merck) and LPS (3.5 μg/mL, Merck) for 2 days of in vitro culture (until d5). On day 5, activated B cells were resuspended in PBS and divided into two fractions, one containing plain PBS and the other additionally supplemented with GP61 peptide (5μM). Cells were kept in the incubator for 1h, after which the fractions were stained with different concentrations of CellTrace Violet (Thermo-Fisher, CTV) (5 μM for GP61-80 loaded cells; 0.5 μM for unloaded cells). B cells were then counted and mixed back together to a 1:1 ratio. The mixed B cell population was eventually added to Th1 CD4^+^ T cells at different ratios ranging from 0X to 16X for 6h in the incubator (Th1 cells : B cells). Cells were then stained with the following markers: CD4□ (RM4-5, eBiosciences), CD19 (ID3, eBiosciences) and Zombie NIR (Biolegend). Samples were immediately acquired at a FACS Canto II after this step.

Calculation of specific killing at every given ratio was done using the following formula:

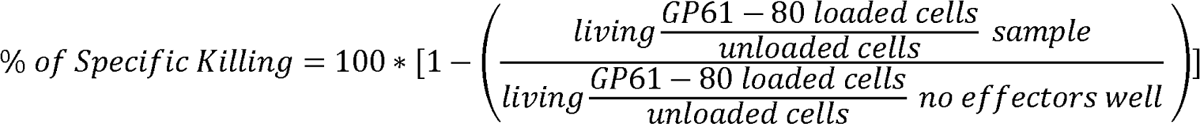

### FACS acquisition and analysis of data

All the samples were acquired in a Canto II cytometer from BD-Biosciences (unless stated otherwise). For the analysis of the data, FlowJo 10.4.2 or posterior versions were used. Regarding the gating strategy, T cells were selected according to their cell size and granularity with the FSC-A and SSC-A, we excluded duplets by SSC-H vs. SSC-W and FSC-H vs. FSC-W gating. Finally, we excluded dead cells by gating on CD4^+^ZombieAqua^-^ cells. For the quantification of frequencies and MFI, statistics were exported to excel files, normalized and transferred to GraphPadPrism for the generation of plots and statistical analysis.

### Seahorse analysis

Samples were washed and resuspended in Agilent Seahorse XF RPMI Medium pH 7.4 (Agilent) containing glutamine (2mM, Gibco), glucose (10mM, Sigma), Na-pyruvate (1mM, Sigma) and lactate when indicated. Cells were counted, adjusted to a final concentration of 1.1x10^6^ cells/mL and seeded in a 96-well utility plate precoated with poly-D-lysine (50μg/mL, Sigma). After a resting period (1h; incubator without CO_2_), cells were placed in a SeaHorse XFe96 analyzer to track OXPHOS (OCR) and glycolytic (PER) activity. After assessment of the steady-state metabolic activity, oligomycin (Oligo, 10.5μM, Sigma), FCCP (7.74μM, Cayman Chemical) and rotenone/antimycin A (R/AA, 11.5μM and 9μM, Cayman Chemical and Sigma, respectively) were injected in a sequential fashion. At least 4 technical replicates were plated for every sample. Initial analysis was done in Wave for bubble exclusion, final analysis was done in GraphPad Prism, where the technical replicates of every sample were averaged and pooled for every biological sample. Basal OCR and PER were defined from the 4th measurement point before oligomycin injection, whereas maximal OCR and PER were defined from the first measurement point after FCCP injection. Metabolic plasticity was calculated for each pathway as the difference between maximal and basal values and is reported as spare respiratory capacity (SRC) for OXPHOS and glycolytic reserve (GR) for glycolysis. For human Seahorse measurements, OCR and PER values of each sample were normalized to the first measurement point of the paired control donor to account for inter-donor variability.

### Isotopic labeling

On day 5, both control and D-lactate-treated cells were plated for 24 hours in the presence of [U-^13^C_3_] D-lactate (20mM; Merck). Extraction of intracellular metabolites, GC-MS, MID calculations and determination of fractional carbon contributions were performed as previously described ^68^ using the Metabolite Detector software package. Glucose, lactate and amino acid concentrations were detected with a YSI 2959D Biochemistry Analyzer (YSI Incorporated).

### RNA isolation, cDNA synthesis and qPCR analysis

RNA was purified and converted to cDNA using a Nucleospin RNA XS kit (Macherey Nagel) and Taqman Reverse Transcription reagent kit (Thermo-Fisher) according to manufacturer’s instructions, respectively. qPCRs were carried out in a Quant Studio 7 Flex Real-time qPCR system (Thermo-Fisher). For every tested gene, 3 technical replicates were generated. The relative expression of every gene was always normalized to the mean value of the house-keeping gene (*Hprt*) on the same sample. The following primer pairs were used: *Ldhd*: Ldhd-fwd (GAATCGATGCACAGGTGTCAACCTC), Ldhd-rev (CAGCTCCGTGATCTGGTCCATAT); *Ifng*: Ifng-fwd (GCCACGGCACAGTCATTGA), Ifng-rev (TGCTGATGGCCTGATTGTCTT) ^69^; *Hprt*: Hprt-fwd (CATAACCTGGTTCATCATCATCGC), Hprt-rev (TCCTCCTCAGACCGCTTTT).

### RNA-Seq library preparation and analysis

Cells were analyzed either under resting conditions or after a short restimulation with plate bound anti-CD3 and anti-CD28 for 4 hours. Isolated RNA quality was assayed with a Fragment Analyzer System (Agilent), and all samples displayed high RNA quality (RQN > 8). cDNA libraries were generated using a Trueseq Stranded Total RNA Library kit (Illumina) according to manufacturer’s instructions. Paired-end sequencing (2x75 bp) was performed on an Illumina NextSeq500 device using a NextSeq500/550 High Output Kit v2. Reads were mapped to the mm10 genome and differential expression was assessed with edgeR and limma^70,71^. Low-expressed genes were filtered and libraries were normalized by TMM^72^. A design matrix modelling all conditions was built, contrasts were defined with makeContrasts, and counts were transformed with voomWithQualityWeights before fitting per-contrast linear models and applying empirical Bayes moderation. Genes were ranked by −log□□(P) × sign(log□FC), with P floored at 10□³□□, and used for GSEA of GO Biological Process terms with clusterProfiler^73^, collapsing redundant terms with simplify. Transcription factor activity was inferred from normalized expression using the univariate linear model in decoupleR^74^ with mouse DoRothEA regulons (confidence levels A–C)^75^, estimating activity from target-gene rather than transcription factor expression.

### ATAC-Seq library preparation and analysis

Libraries were prepared using an ATAC-Seq Kit (Active Motif) according to manufacturer’s instructions and paired-end sequenced (2x76 nt) on an Illumina NextSeq2000 device after quality control on a Fragment Analyzer System and quantification with Kapa and Qubit assays.

Reads from control, D-lactate and L-lactate conditions (acute and prolonged) were quality-checked and trimmed with FastQC and fastp^76^, aligned to mm39 with Bowtie2^77^ (v2.4.4), and filtered with SAMtools (v1.23)^78^ to remove low-quality reads (MAPQ < 30), duplicates and blacklisted regions. Peaks were called with LanceOtron^79^ and merged into a consensus set, and a count matrix was generated with featureCounts^80^ (Subread v2.1.1). Differential accessibility was assessed with edgeR and limma as for RNA-seq; significant regions were annotated with ChIPseeker^81^ and subjected to GO enrichment with clusterProfiler^73^. Motif enrichment was performed with HOMER^82^, and transcription factor footprinting with TOBIAS (v0.13.3)^40^. RNA-seq and ATAC-seq were aligned to mm10 and mm39, respectively; the two datasets were not integrated at the level of genomic coordinates.

## QUANTIFICATION AND STATISTICAL ANALYSIS

To pool results from different experiments, data belonging to the same experiment were normalized to the mean of the control group (Th1 without lactate) and pooled in GraphPad Prism. Unless stated otherwise, data are presented as mean ± SD and each symbol represents an individual biological replicate of normalized pooled data. The number of independent experiments contributing to each panel was: Figure 1, 17 (B, IFN-γ), 15 (B, TNF-α), 8 (F, RUNX3 d5), 5 (E and F, RUNX3 d10), 4 (F, EOMES) and 2 (C, H); Figure 2, 3 (B, D for IFN-γ) and 2 (C, D for TNF-α); Figure 4, 3 (C–E); Figure 5, 6 (A), 3 (C) and 1 (B, F); Figure 6, 5 (E, G), 4 (B) and 3 (C); Figure 7, 6 (B), 4 (H, I) and 3 (D, E). RNA-seq and ATAC-seq analyses (Figures 2E–2H, 3, 4A, 4B, 5D and 6A) were performed with two biological replicates per condition.

Where group size was sufficient to support it, distributions were assessed for normality using the normality tests implemented in GraphPad Prism; parametric tests were applied to normally distributed data and non-parametric tests otherwise. All analyses were performed in GraphPad Prism. The statistical test applied to each panel, together with any correction for multiple comparisons, is indicated in the corresponding figure legend. Significance is indicated as *p < 0.05; **p < 0.01; ***p < 0.001; ****p < 0.0001. Exact p values are given for selected comparisons that did not reach significance.

## Acknowledgments

We thank N.A. Durán-Hernández, I. Panse, V. Holecska, and C. Rüster for expert technical assistance and M. Villa, D. Zaiss and G. Krönke for helpful scientific discussions. We additionally thank P. Durek and F. Heinrich for their help uploading the ATAC-seq and RNA-seq samples to public repositories. Cell sorting was carried out at the Flow Cytometry Core Facility of the DRFZ.

## Funding

This work was supported by the Deutsche Forschungsgemeinschaft (DFG; grants LO 1542/4-1 and LO 1542/5-1 to M.L.), the Einstein Center for Regenerative Therapies (ECRT; grant EZ-2016-289 to A.M.A. and V.P.), the Willy Robert Pitzer Foundation (Pitzer Laboratory of Osteoarthritis Research; grant 21-033 to M.L.), and the Dr. Rolf M. Schwiete Foundation (Osteoarthritis Research Program; grant 2021-035 to M.L.). D.B. and L.S.B. were supported by the Luxembourg National Research Fund (FNR; grants C21/BM/15796788 and PRIDE/11012546/NEXTIMMUNE).

## Author contributions

Conceptualization: AMA, ML

Methodology: AMA, PRVM, ML

Investigation: AMA, VP, LSB

Formal analysis: AMA, PRVM, SS, MD

Resources: DB, MFM

Visualization: AMA

Project administration: AMA

Supervision: AMA, ML

Funding acquisition: AMA, VP, DB, ML

Writing—original draft: AMA

Writing—editing: AMA, VP, PRVM, LSB, DB, MD, ML

Writing—reviewing: All authors

## Declaration of interests

The authors declare no competing interests.

---

**Figure S1.**
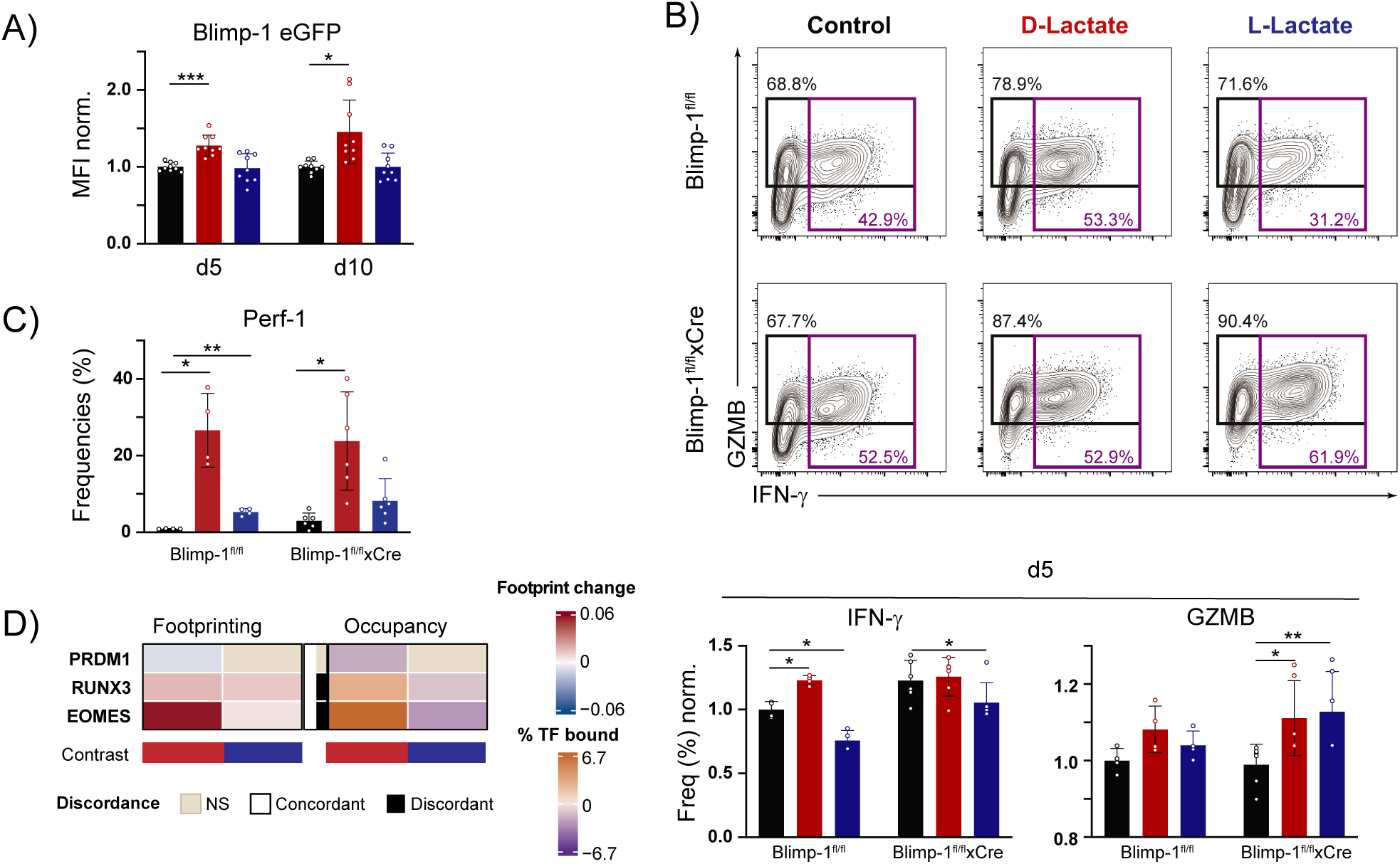
**D-lactate uncouples Blimp-1 induction from repression of IFN-γ production in Th1 cells**. (A-D) Sorted naïve mouse WT, Blimp-1eGFP, Blimp-1^fl/fl^ and Blimp-1^fl/fl^ xCre CD4^+^ T cells were cultured for 5 (A, B, D) or 10 (C) days under Th1-polarizing conditions with or without lactate supplementation, as indicated. (A) Quantification of Blimp-1 expression levels (n=9). (B) Representative contour plots (top) and quantification (bottom) of IFN-γ and GZMB production in Blimp-1^fl/fl^ and Blimp-1^fl/fl^ xCre Th1 cells on day 5 (n=4, Blimp-1^fl/fl^; n=6 Blimp-1^fl/fl^ xCre). (C) Quantification of Perf-1 expression in Blimp-1^fl/fl^ and Blimp-1^fl/fl^ xCre Th1 cells on day 10 (n=4, Blimp-1^fl/fl^; n=6 Blimp-1^fl/fl^ xCre). (D) Heatmaps representing footprint (left) and occupancy (right) changes for PRDM1, RUNX3 and EOMES across the indicated contrasts, derived from ATAC-seq data generated from two biological replicates per condition. Two-way ANOVA (A) or Mixed-effects (B,C), both with Dunnett’s post-tests.

**Figure S2.**
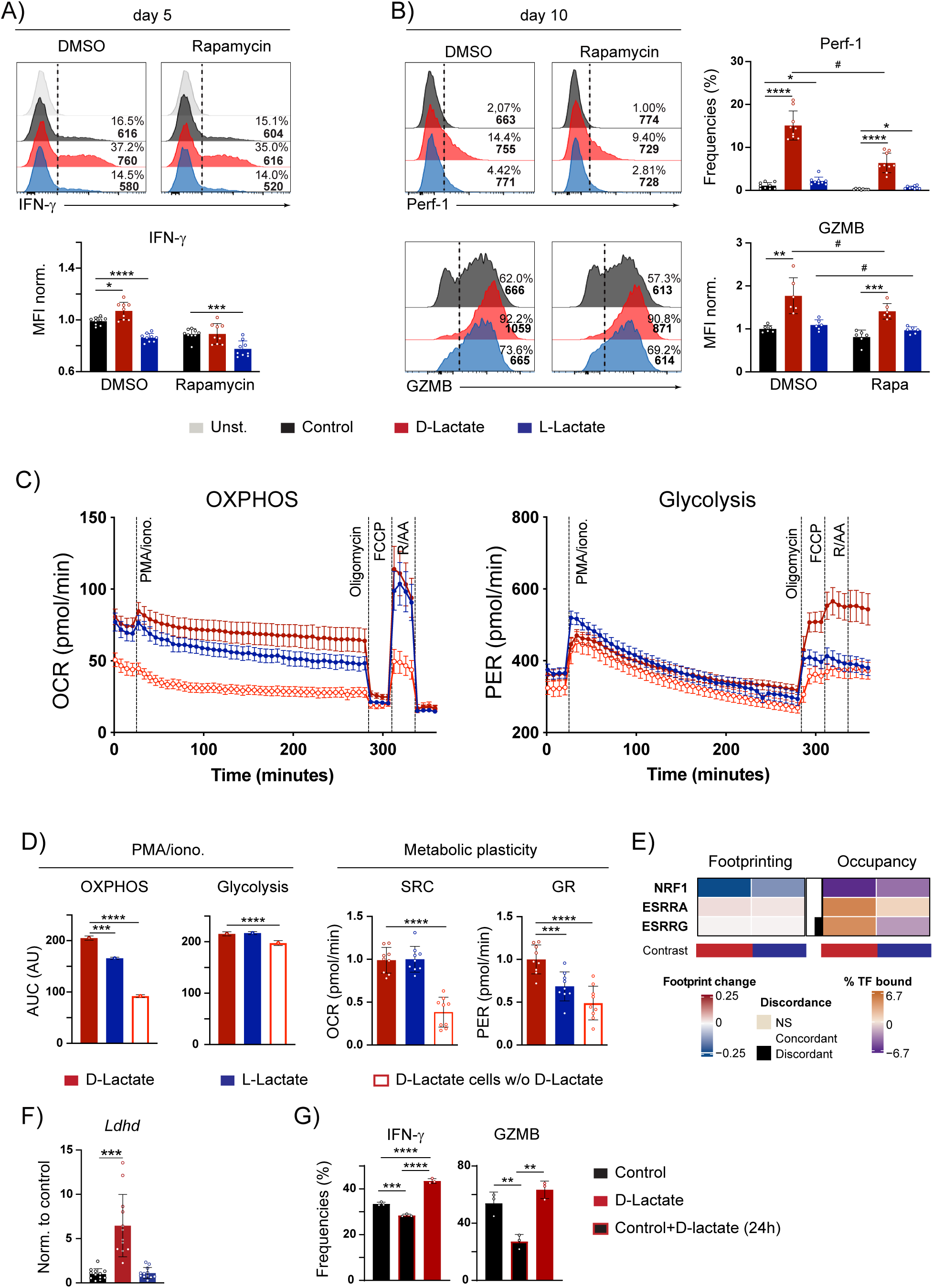
**mTORC1 contributes to D-lactate-driven effector features, whereas acute exposure induces a distinct metabolic and functional state**. (A-G) Sorted naïve WT CD4^+^ T cells were cultured for 5 (A, C-F), 6 (G) or 10 (B) days under Th1-polarizing conditions with or without lactate supplementation as indicated. (G) On day 5, control and D-lactate-treated cells were replated in RPMI medium containing [U-13C] D-lactate for 24 hours. (A) Representative histograms (top) and quantification (bottom) of IFN-γ production after 4h of rapamycin (100nM) treatment on day 5 (n=9). (B) Representative histograms (left) and quantification (right) of Perf-1 (top) and GZMB (bottom) production after 4h of rapamycin treatment on day 10 (n=9). (C) Representative Seahorse kinetics for OXPHOS (left) and glycolysis (right) (n=9). (D) Quantification of OXPHOS and glycolytic activity during the 4 h PMA/ionomycin restimulation period (left), together with their corresponding metabolic plasticity (right) (n = 9). (E) Heatmaps representing footprint (left) and occupancy (right) changes for NRF1, ESRRA and ESRRG across the indicated contrasts, derived from ATAC-seq data generated from two biological replicates per condition. (F) qPCR analysis of *Ldhd* expression (n = 12). (G) Quantification of IFN-γ and GZMB production after 24h of [U-^13^C_3_] D-lactate treatment (n=3). Two-way ANOVA with Dunnett’s post-tests (A and B) or One-way ANOVA with either Tukey’s (G) or Dunnett’s (D, F) post-tests.

**Figure S3.**
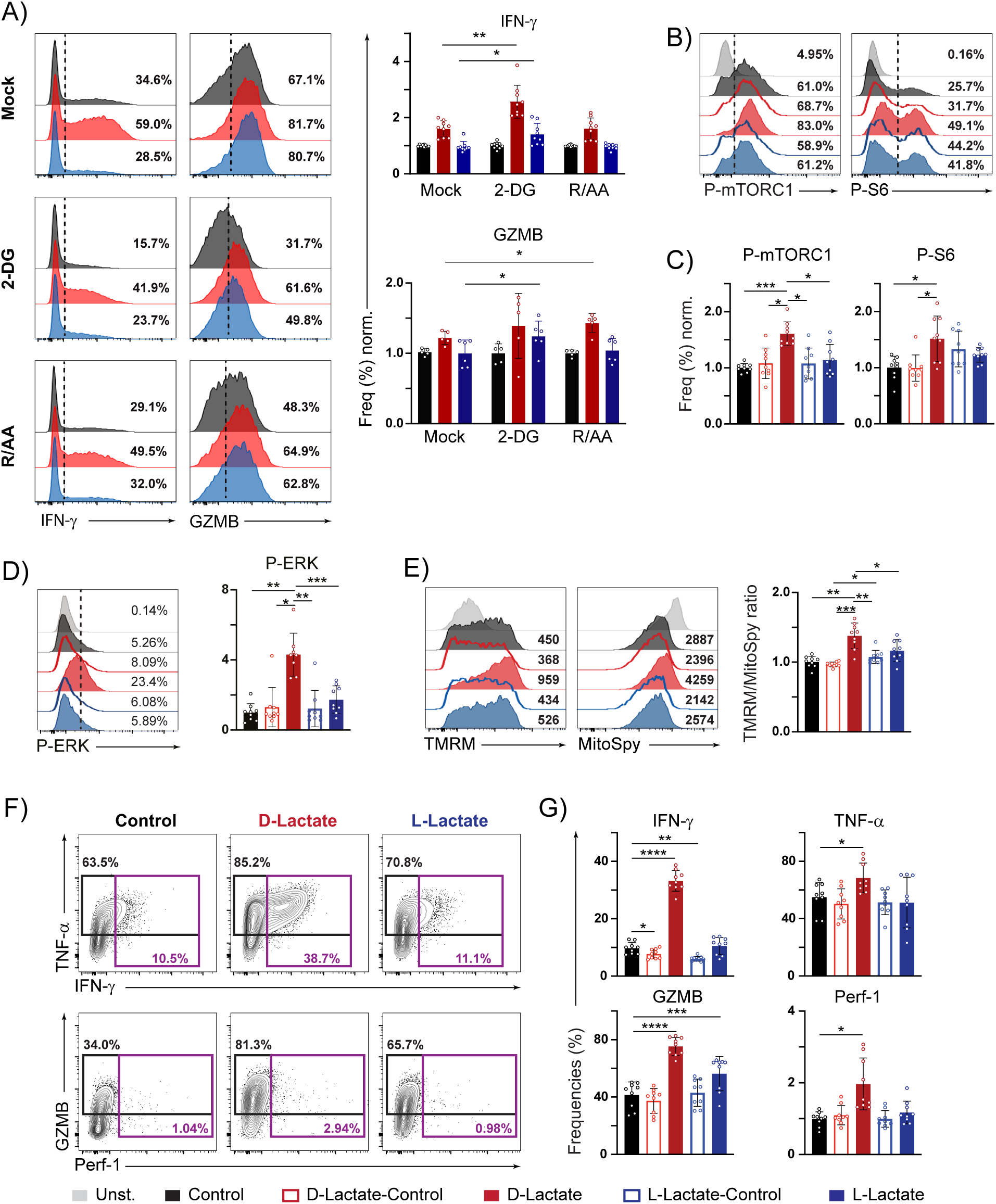
**D-lactate sustains Th1 effector function under glucose-restricted conditions**. (A-G) Sorted naïve mouse WT CD4^+^ T cells were cultured for 5 days under Th1-polarizing conditions with or without lactate supplementation, as indicated (A). On day 5 cells were replated in glucose-restricted medium (1 mM) for three additional days, a fraction of the lactate-differentiated samples was cultured in absence of exogenous lactate supplementation (B-G). (A) Representative histograms (left) and quantification (right) of IFN-γ and GZMB production after 4 hours of either 2-DG (10 mM) or R/AA (11.5 μM and 9 μM) treatment (n=6). (B) Representative histograms of P-mTORC1 (S2448) (left) and P-S6 (S240/S244) (right). (C) Quantification of (B) (n=9). (D) Representative histogram (left) and quantification (right) of P-ERK (n=9). (E) Representative histograms (left) and quantification (right) of TMRM and MitoSpy Green staining (n=9). (F) Representative contour plots of TNF-α, IFN-γ (top) and GZMB, Perf-1 (bottom). (G) Quantification of (F) (n=9). Two-way ANOVA with Dunnett’s post-tests (A) or RM-One-way ANOVA with either Tukey’s (C-E) or Dunnett’s (G) post-tests.

**Figure S4.**
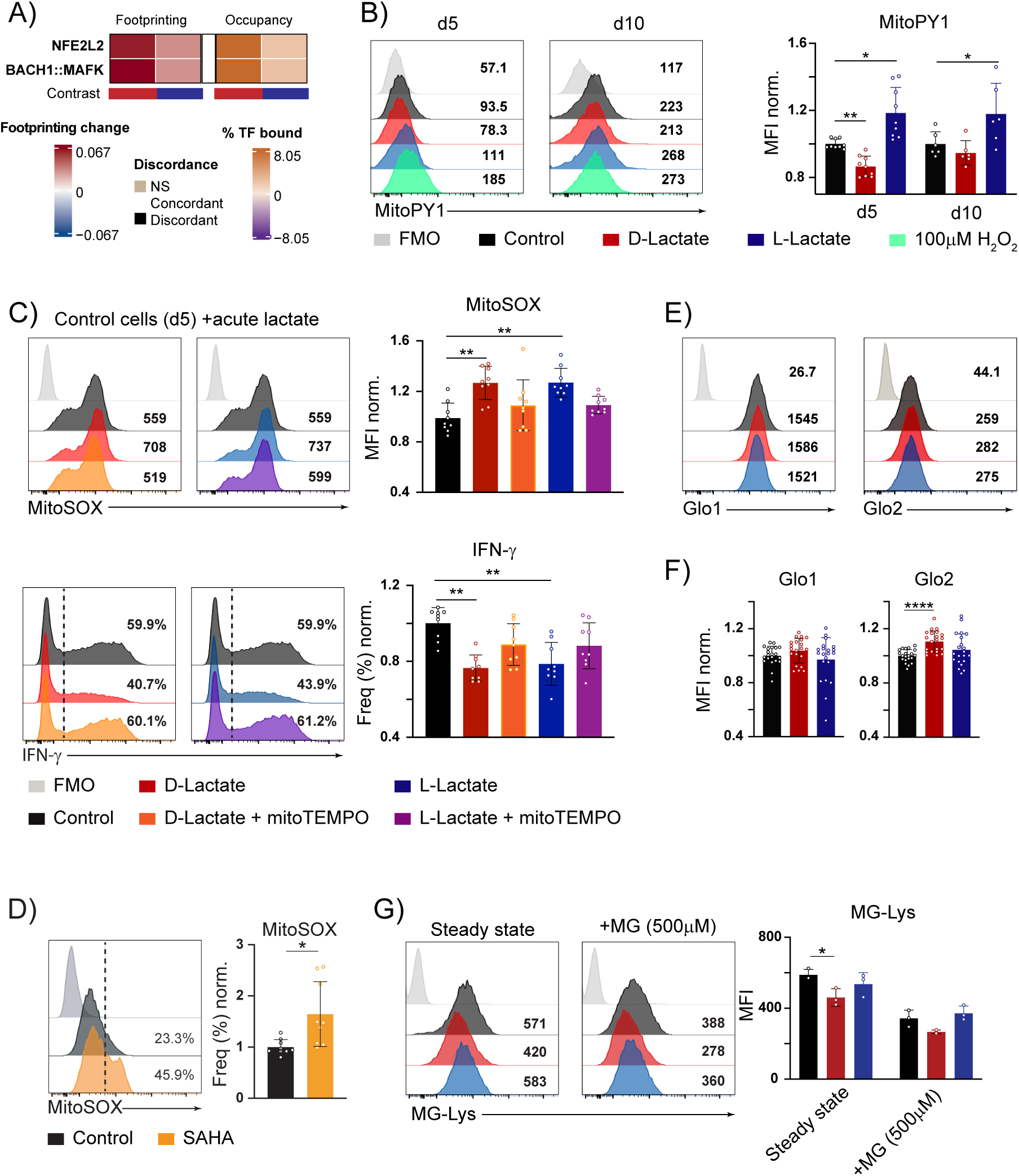
**D-lactate promotes a distinct redox state associated with the glyoxalase system**. (A-G) Sorted naïve mouse WT CD4^+^ T cells were cultured under Th1-polarizing conditions for 5 days with or without lactate supplementation, as indicated. (A) Heatmaps representing footprint (left) and occupancy (right) changes for NFE2L2 and BACH1 across the indicated contrasts, derived from ATAC-seq data generated from two biological replicates per condition. (B) Representative histograms (left) of MitoPY1 staining and quantification(right) (n=9). (C) Representative histograms (left) and quantification (right) of MitoSOX (top) and IFN-γ (bottom) staining (n=9). (D) Representative histogram (left) and quantification (right) of MitoSOX staining after 4h of SAHA treatment during PMA/ionomycin restimulation (n=9). (E) Representative histograms of Glo1 and Glo2 staining. (F) Quantification of (E) (n=18). (G) Representative histograms (left) and quantification (right) of MG-Lys staining (n=3). Two-way ANOVA with Dunnett’s post-tests (B and G), RM-One-way ANOVA with either Dunnett’s (C) or Tukey’s post-tests (F) or two-tailed t tests (D).

## Notes

### Competing Interest Statement

The authors have declared no competing interest.

### Summary of Updates

This version has been revised with an updated introduction, revised figures and figure legends, and updated author information. The text was edited throughout for clarity.

